# Generation of circulating autoreactive pre-plasma cells fueled by naïve B cells in celiac disease

**DOI:** 10.1101/2023.10.20.563227

**Authors:** Ida Lindeman, Lene S. Høydahl, Asbjørn Christophersen, Louise F. Risnes, Jørgen Jahnsen, Knut E. A. Lundin, Ludvig M. Sollid, Rasmus Iversen

## Abstract

Autoantibodies against the enzyme transglutaminase 2 (TG2) are characteristic of celiac disease (CeD), and TG2-specific IgA plasma cells are abundant in gut biopsies of patients. We here describe the corresponding population of autoreactive B cells in blood. Circulating TG2-specific IgA cells were found in untreated patients on a gluten-containing diet but not in controls. They were clonally related to TG2-specific small intestinal plasma cells, and they expressed gut-homing molecules, indicating that they are plasma cell precursors. Unlike other IgA-switched cells, the TG2-specific cells were negative for CD27, placing them in the double negative (IgD-CD27-) category. They had a plasmablast or activated memory B-cell phenotype, and they harbored fewer variable region mutations than other IgA cells. Based on their similarity to naïve B cells, we propose that autoreactive IgA cells in CeD are generated mainly through chronic recruitment of naïve B cells via an extrafollicular response involving gluten-specific CD4+ T cells.

## INTRODUCTION

Autoantibody production against the enzyme transglutaminase 2 (TG2) is tightly linked to celiac disease (CeD). Both IgA- and IgG-class autoantibodies can be detected in patient blood samples, but testing for anti-TG2 IgA shows the highest diagnostic accuracy.^1^ In addition, there is accumulation of autoreactive plasma cells in the small intestinal mucosa with anti-TG2-producing cells typically making up 10-20% of all the lamina propria plasma cells in patients.^2,3^ Most of these cells secrete IgA antibodies, and they show a clonal relationship with TG2-specific IgA in serum.^4^ Both the serum antibodies and the gut plasma cells are strictly gluten dependent, as serum anti-TG2 levels drop and TG2-specific plasma cell frequencies decrease within a few months after patients commence a gluten-free diet.^5,6^

The gluten dependence of the autoantibody response can be explained by T cell-B cell collaboration involving gluten-specific CD4+ T cells and TG2-specific B cells.^7^ In CeD, T cells specifically recognize deamidated gluten peptides in which certain glutamine residues have been converted to glutamic acid. This deamidation happens in the human body and is catalyzed by TG2.^8,9^ Since many gluten peptides are substrates for the enzyme, TG2-specific B cells can take up TG2-gluten enzyme-substrate complexes via their B-cell receptor (BCR) followed by presentation of deamidated gluten to T cells.^10^

Most of the TG2-specific plasma cells in CeD belong to the short-lived CD19+ subset, and they generally harbor fewer immunoglobulin (Ig) mutations than other gut plasma cells.^3,11^ Since generation of long-lived plasma cells and accumulation of high levels of Ig mutations are typical features of germinal center (GC) reactions,^12^ these observations may suggest that autoantibody formation in CeD depends on extrafollicular B-cell activation. However, the B-lineage cells that can be extracted from gut biopsies are terminally differentiated tissue-resident plasma cells, and they do not provide direct insight into earlier stages of B-cell development. To better understand the mechanisms of autoantibody formation in CeD, we here aimed to identify circulating TG2-specific B cells in patient blood samples, as such cells presumably represent the precursor population of the gut plasma cells.

Our results show that IgA-switched TG2-specific B cells are uniquely present in untreated CeD patients. These cells have a gut-homing, activated phenotype, and they are clonally related to tissue-resident gut plasma cells. Curiously, most of the cells do not express the classical memory B-cell marker CD27. They hence belong to the double negative (DN; IgD-CD27-) fraction of B cells.^13^ However, they do not fall in the so-called DN2 category, which has previously been associated with autoimmunity and some chronic infectious diseases.^14,15^ Rather, the cells appear to represent an early stage of B-cell differentiation resulting from chronic, gluten-dependent immune reactions in which naïve B cells are continuously recruited and pushed into a plasma cell differentiation pathway.

## RESULTS

### TG2-specific IgA B cells in untreated CeD lack expression of CD27

We sought to detect circulating autoreactive B cells in CeD by staining patient blood samples with recombinant human TG2 (Figure S1A). In addition, glutathione S-transferase (GST) was included as an irrelevant control antigen to increase staining specificity. Flow cytometry analysis demonstrated that TG2-specific cells made up a small proportion of IgA+ B cells in blood (mean 0.4%, range 0.3-1.0%, Figure 1A). However, the frequency could be increased around 20-fold by including an antigen-specific enrichment step prior to analysis. When considering all enriched B cells, a TG2-specific population could be detected in both untreated patients and treated patients on a gluten-free diet as well as in healthy donors (Figure 1B). In all cases, naïve (IgD+CD27-) B cells made up most of the TG2-specific cells. Only in CeD patients, however, did we detect a population of IgA-switched autoreactive cells. These cells were present in untreated patients, but numbers were greatly reduced in most patients who had been on a gluten-free diet for at least one year (Figures 1B and 1C). Interestingly, two treated patients who had high levels of circulating TG2-specific IgA cells both showed incomplete mucosal recovery, possibly due to poor adherence to gluten-free diet (Figure 1C). Unlike other IgA+ B cells, the TG2-specific cells in untreated patients were predominantly negative for the classical memory B-cell marker CD27 (Figures S1A and 1D). Lack of CD27 expression correlated with the frequency of TG2-specific IgA cells (Figure 1E), suggesting that the CD27-cells result from ongoing immune reactions in active CeD, where high numbers of autoreactive B cells are generated. Since these DN cells were largely absent in patients on a gluten-free diet, they do not appear to persist in the blood as long-lived memory B cells.

**Figure 1.**
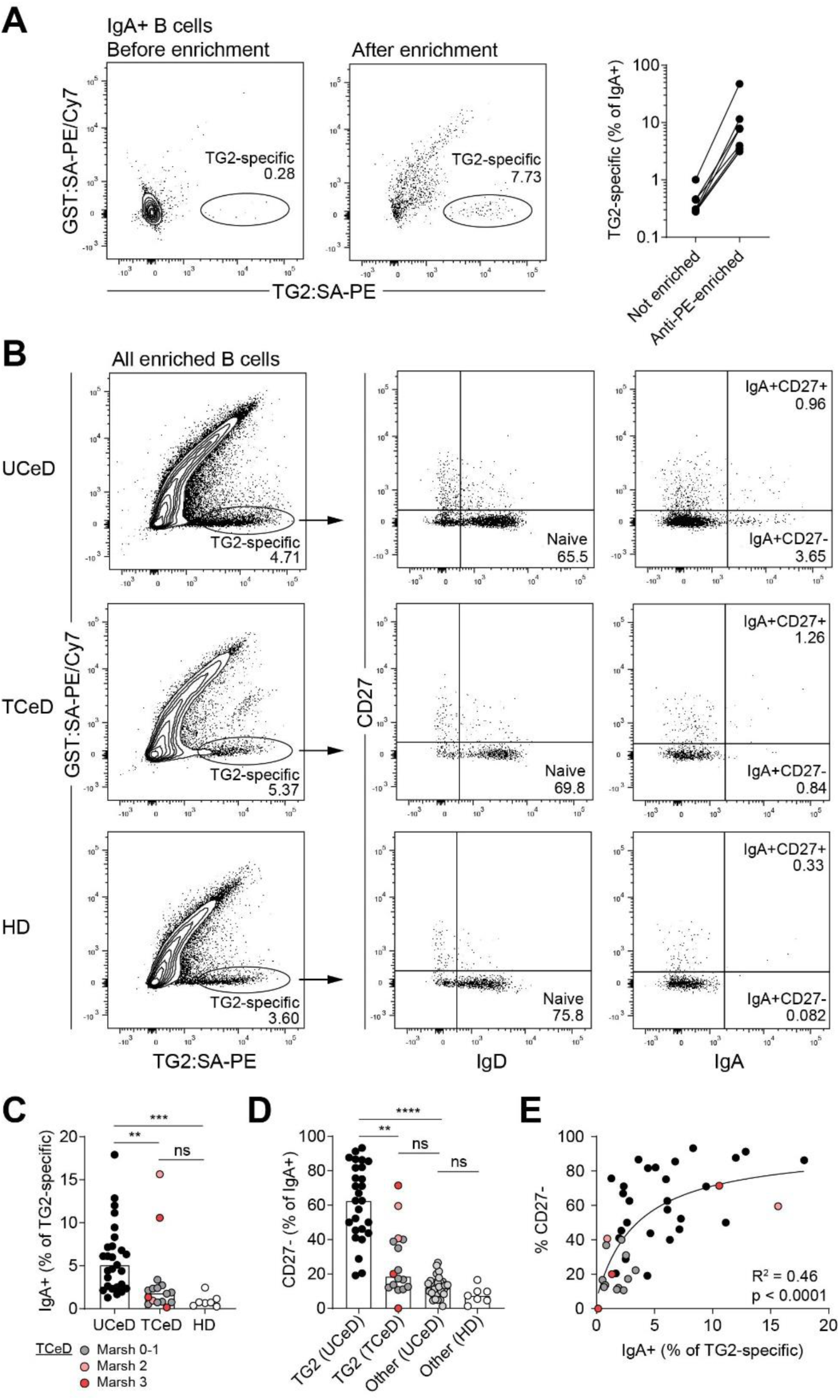
Identification of circulating TG2-specific B cells. (A) Representative flow cytometry plots and summary data from eight CeD patients showing TG2 staining of IgA-switched B cells in peripheral blood before and after antigen-specific enrichment. GST was included as an irrelevant control antigen. The biotinylated antigens were attached to streptavidin (SA) that was conjugated to PE or PE/Cy7, allowing enrichment with anti-PE microbeads. (B) Representative flow cytometry plots showing identification of TG2-specific cells among all B cells after antigen-specific enrichment. Blood samples were obtained from patients with untreated CeD (UCeD), treated CeD (TCeD) or from healthy donors (HD). (C) Frequency of IgA-switched cells among all TG2-specific B cells in UCeD patients (n=27), TCeD patients (n=16) or HD (n=7) after antigen-specific enrichment. Mucosal healing in TCeD was evaluated by Marsh score classification, where 0 or 1 is considered normal. (D) Frequency of CD27-cells among TG2-specific and non-TG2-specific (other) IgA+ B cells in the same donors. Bar heights indicate medians, and difference between groups was evaluated by a Kruskal-Wallis H test with Dunn’s multiple comparisons correction. **p < 0.01, ***p < 0.001, ****p < 0.0001. (E) Correlation between frequency of IgA-switched cells and frequency of CD27- IgA-switched cells among TG2-specific B cells. Symbol colors indicate UCeD or TCeD as in (C). The data were fitted to a three-parameter dose-response curve, and the p value indicates Spearman correlation.

### Autoreactive B cells are functional and secrete TG2-specific IgA in vitro

To confirm that the autoreactive cells we could detect by flow cytometry were truly TG2-specific, we established a culture system for human B cells based on what has previously been used for in vitro activation of mouse B cells (Figures S2A-S2C).^16,17^ Single TG2-specific or non-TG2-specific IgA+ B cells were sorted from patient blood samples and placed in culture with transfected NIH/3T3 fibroblasts expressing human CD40 ligand (CD40L), human B-cell activating factor (BAFF) and human IL-21. After nine days, antibody secretion from the single-cell cultures was detected by ELISA. Most of the cells that bound to TG2 in flow cytometry and secreted detectable levels of IgA in vitro also gave rise to an anti-TG2 signal, thus confirming their specificity (Figures 2A and S2D). There was no difference in total IgA secretion between TG2-specific and non-TG2-specific B cells, nor was there a difference between CD27+ and CD27- cells (Figure 2B). Most of the sorted autoreactive B cells were negative for CD27, and these were more often confirmed to be TG2-specific after culturing than the more rare CD27+ cells (Figure 2C). However, among the cells with confirmed TG2-specificity there was no apparent difference in affinity between CD27+ and CD27- cells (Figure 2D). Collectively, our results show that the autoreactive B cells we detect in patient blood samples are functional and capable of secreting high levels of TG2-specific IgA after in vitro activation.

**Figure 2.**
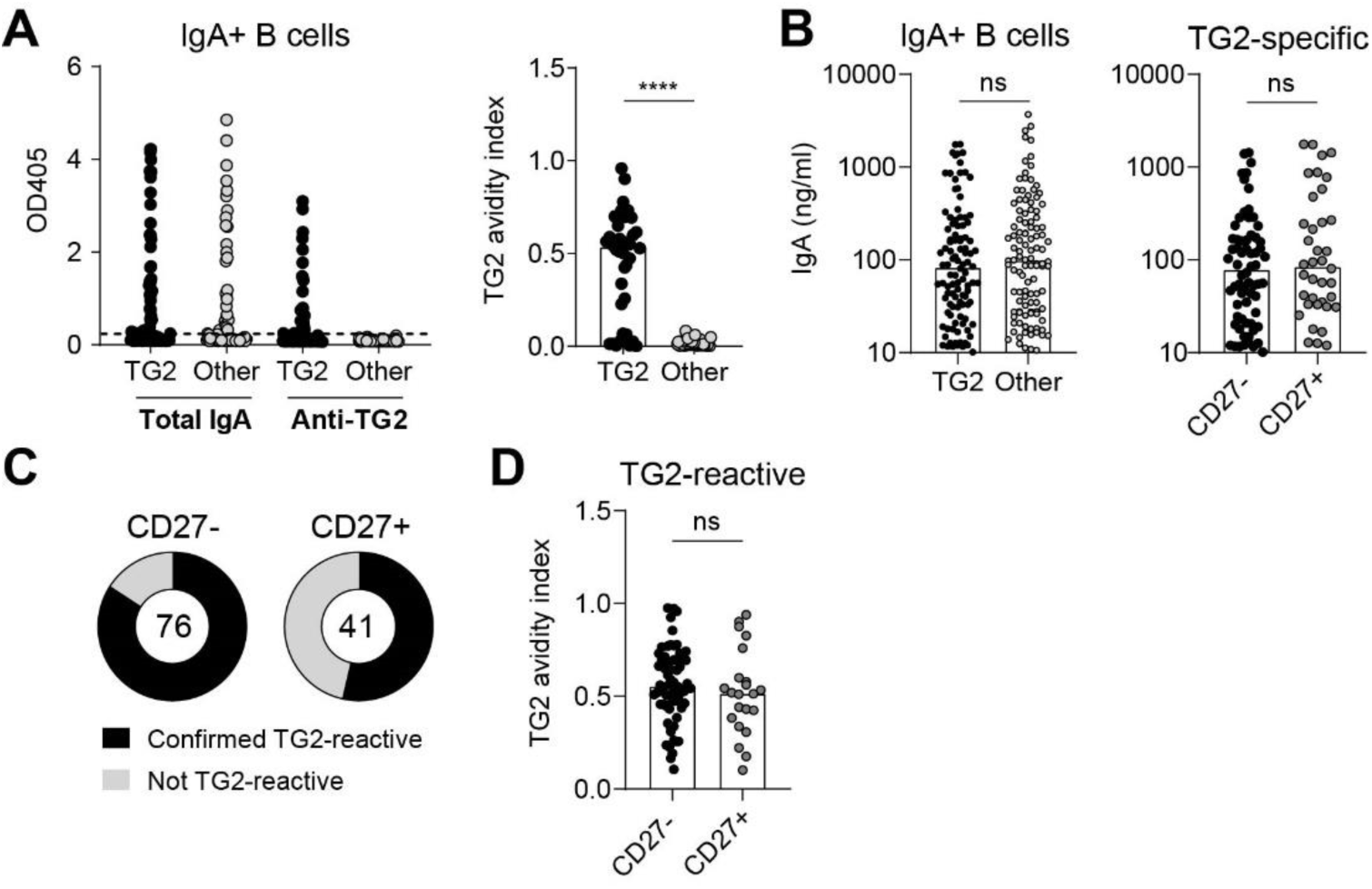
IgA secretion in single-B-cell cultures. (A) Detection of total and anti-TG2 IgA in culture supernatants of single TG2- specific (n=88) or other (n=88) IgA+ B cells sorted from PBMCs of a representative untreated CeD patient. The TG2 avidity index was calculated as the TG2-specific signal divided by the total IgA signal for cultures positive for total IgA (OD > 3x background, dashed line). (B) Comparison of antibody secretion (>10 ng/ml total IgA) in cultures of TG2-specific (n=106) or other (n=111) IgA+ B cells sorted from PBMCs of four untreated CeD patients. (C) Confirmation of TG2-reactivity in all cultures of TG2-specific B cells that were positive for total IgA. A TG2 avidity index >0.1 was considered TG2 positive. (D) Affinity of secreted antibodies in cultures of CD27+ or CD27- IgA+ B cells with confirmed TG2 reactivity. Bar heights indicate medians, and difference between groups was evaluated by a Mann-Whitney U test. ****p < 0.0001.

### TG2-specific gut plasma cells in CeD are negative for CD27

To investigate if the phenotype of circulating TG2-specific B cells would be mirrored by tissue-resident gut plasma cells, we first analyzed duodenal biopsies of untreated CeD patients by flow cytometry (Figure S1B). Although release of endogenous TG2 during tissue processing in combination with high levels of ambient TG2-specific IgA cause some TG2-specific IgM plasma cells to appear as surface IgA+, most of the TG2-binding IgA+ cells in gut biopsies are true IgA plasma cells (Figure S3). As seen in blood, most of the TG2-specific cells were negative for CD27 (Figures 3A and 3B). Further, the CD27- cells were enriched for CD19+ short-lived plasma cells (Figures 3A and 3C). These observations agree with previously reported bulk and single-cell RNA sequencing (scRNA-seq) data showing that TG2- specific plasma cells have lower expression of *CD27* than other gut plasma cells^3,18^ and that low *CD27* expression is associated with a short-lived (CD19+CD45+) rather than intermediate- (CD19-CD45+) or long-lived (CD19-CD45-) plasma cell phenotype.^3^ To gain further insight into potential phenotypic differences between CD27+ and CD27- plasma cells, we next performed droplet-based scRNA-seq in combination with CITE-seq,^19^ thereby allowing simultaneous transcriptomic analysis and detection of surface CD27 and TG2-specific BCRs (Figure 3D and Table S1). V(D)J sequence analysis of TG2-specific and non-TG2-specific IgA plasma cells revealed that individual clonotypes often included both CD27+ and CD27- cells (Figures 3E and 3F). Further, CD27- cells were interspersed between CD27+ cells in clonal lineage trees, and in some cases, identical sequences were obtained from both types of cells (Figure 3G). These results indicate that CD27 status is not imprinted during initial activation of naïve B-cells. Rather, CD27 expression can be acquired, or potentially lost, as B cells undergo clonal diversification. When comparing gene expression between surface CD27+ and CD27- cells, *CD27* itself stood out as the one gene with clear differential expression (Figure 3H). Thus, CD27- plasma cells appear functionally similar to CD27+ plasma cells at the transcriptional level. Among the few other genes found to have higher expression in CD27+ cells, *S100A6* was previously observed to be upregulated in long-lived compared to short-lived plasma cells,^3^ consistent with the observation that CD27- cells primarily belong to the short-lived population.

**Figure 3.**
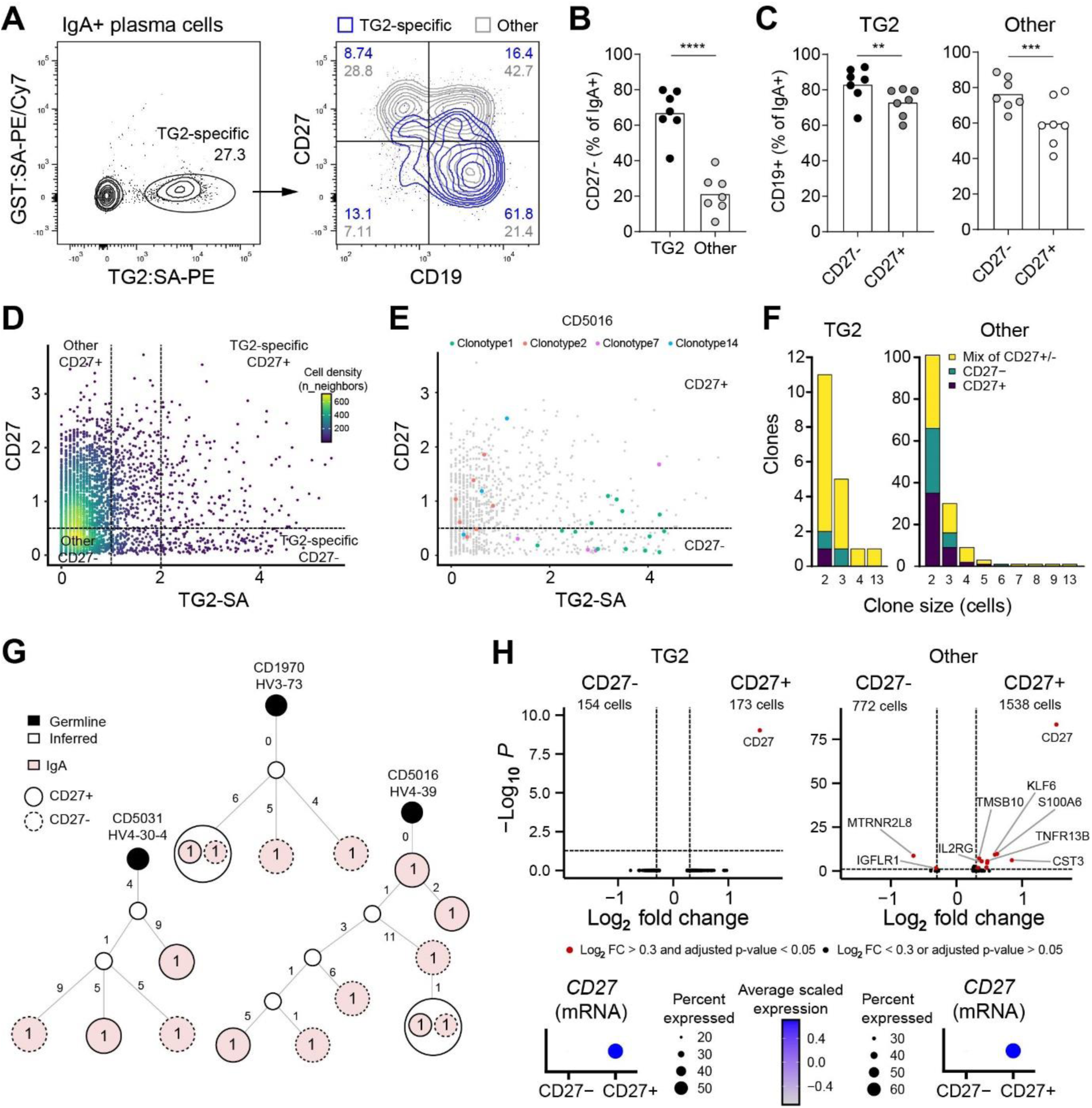
Characterization of CD27+ and CD27- gut plasma cells. (A-C) Representative flow cytometry plot (A) and summary data from seven untreated CeD patients showing staining of duodenal IgA+ plasma cells. The frequency of CD27- (B) and CD19+ (C) cells was evaluated among TG2-specific and non-TG2-specific (other) plasma cells. Bar heights indicate means, and difference between groups was evaluated by a paired t test. **p < 0.01, ***p < 0.001. ****p < 0.0001. (D-H) CITE-seq, V(D)J and transcriptome analysis of IgA+ plasma cells sorted from duodenal biopsies of four untreated CeD patients. Only cells with confirmed expression of a V(D)J sequence in connection with an IgA constant region were included in the analyses. (D) Cumulated CITE-seq data showing identification of TG2-binding and non-TG2-binding populations with and without surface CD27. (E) Examples of individual clonotypes spanning surface CD27+ and CD27- populations in a representative patient. (F) Number of expanded clonotypes consisting of only CD27+, only CD27- or both CD27+ and CD27- plasma cells as a function of the number of cells in each clone. (G) Examples of lineage trees showing TG2-specific clones comprising both CD27+ and CD27- cells. Colored circles represent IGHV sequences observed in individual cells, and numbers next to edges indicate mutations (nt). (H) Volcano plots showing differentially expressed genes between surface CD27+ and CD27- populations (upper panels) and verification of CD27 expression at the mRNA level (lower panels).

### Circulating TG2-specific cells comprise plasmablasts and memory B cells

In some patients, we observed that circulating TG2-specific IgA cells had a typical plasmablast phenotype characterized by high expression of CD38 in combination with downregulation of B-cell markers such as CD20, CD21 and CD24 (Figure 4A). In addition, these cells showed low expression of CXCR5 and high expression of integrin β7, indicating that they are migrating to the intestine rather than circulating through lymphoid tissues. Again, we saw that the autoreactive cells lacked expression of CD27 (Figures 4A and 4B). The frequency of CD38+ TG2-specific cells varied between patients, but they were consistently negative for CD27, making them different from other CD38+ IgA cells (Figure 4C). Since plasmablasts are usually defined by high expression of both CD38 and CD27, we sought to address if these DN cells were true plasmablasts. Sorting of CD27-CD38+ B cells followed by IgA- specific ELISPOT confirmed that at least some of the cells spontaneously secrete antibodies and thus can be considered bona fide plasmablasts (Figure 4D).

**Figure 4.**
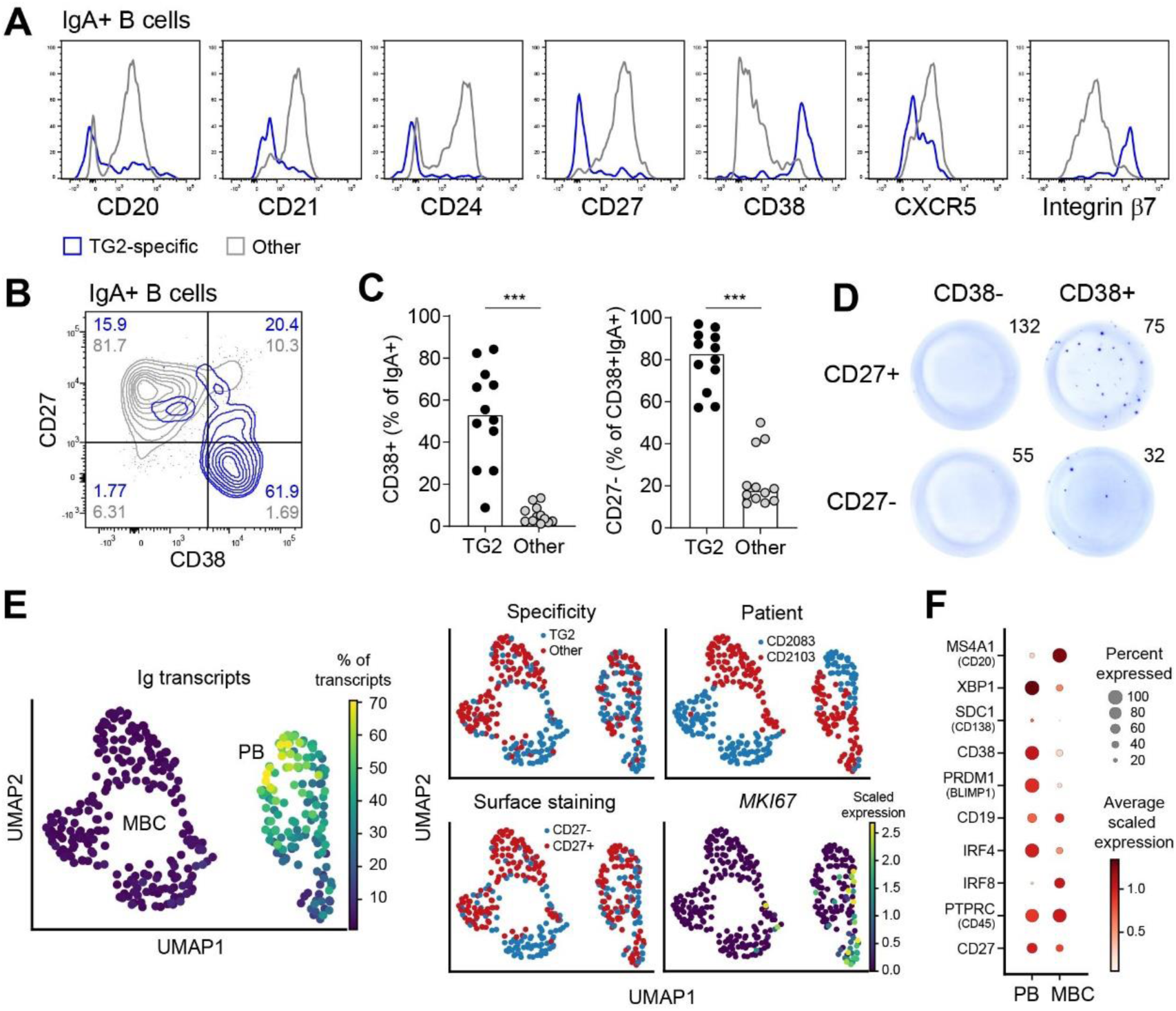
Phenotype of circulating TG2-specific B cells. (A) Representative flow cytometry histograms comparing expression of surface markers associated with a memory B cell or plasmablast phenotype on TG2-specific and non-TG2-specific (other) IgA+ B cells in peripheral blood of an untreated CeD patient. (B and C) Representative flow cytometry plot (B) and summary data from 12 patients (C) showing expression of CD38 and CD27 on IgA+ B cells in blood. Bar heights represent medians, and difference between groups was evaluated by a Wilcoxon signed rank test. ***p < 0.001. (D) ELISPOT detection of IgA- secreting cells among circulating IgA+ B cells with the indicated combinations of surface CD27 and CD38. Numbers indicate the number of sorted cells that were placed in culture in the individual wells. (E) UMAP plot based on scRNA-seq data obtained from TG2-specific and other IgA+ B cells sorted from PBMCs of two untreated CeD patients. Based on gene expression and amount of Ig transcripts, two main clusters were assigned as memory B cells (MBC) and plasmablasts (PB). (F) Expression of genes typically associated with B-cell differentiation in the two clusters.

To gain further insight into the phenotype(s) of circulating autoreactive cells, we next analyzed TG2- specific and non-TG2-specific IgA cells by scRNA-seq using Smart-seq2.^20^ We obtained data from 355 cells isolated from two untreated CeD patients (Table S1). Dimensionality reduction revealed two main cell clusters that we interpret as plasmablasts and memory B cells based on expression of genes typically associated with the two cell types (Figures 4E and 4F). Further, the plasmablast cluster was characterized by high levels of Ig transcripts consistent with antibody production, and a subset of the cells showed expression of *Ki-67*, indicating active proliferation. Notably, the TG2-specific cells were present in both clusters, confirming that circulating autoreactive IgA cells in CeD comprise both plasmablasts and memory B cells.

### Clonal overlap between blood and gut B-cell populations

By reconstructing complete V(D)J sequences from our scRNA-seq data and by sequencing single-B-cell cultures, we obtained the rearranged Ig sequences of individual TG2-specific cells in blood. The cells displayed typical features previously associated with TG2-specific autoantibodies, including preferential usage of particular variable region gene segments and a bias toward the IgA1 subclass (Figures S4A and S4B).^3,4^ There was no obvious connection between V-gene usage and CD27 expression, suggesting that B-cell phenotype is not directly related to targeting of individual TG2 epitopes (Figure S4C).^21^ As seen for gut plasma cells, clonal families in blood comprised both CD27+ and CD27- cells (Figure 5A). In addition, plasmablasts and memory B cells belonged to the same clonotypes. Since we obtained V(D)J sequences from TG2-specific IgA cells in paired samples of blood and duodenal biopsies, we were also able to address the degree of clonal overlap between the two compartments. In samples with sufficient numbers of antigen-specific cells, we observed substantial clonal overlap between B-cell populations in blood and gut, suggesting that relatively few clonotypes make up the majority of the autoreactive cells (Figure 5B). Clonal families spanning blood and gut often contained both plasmablasts and memory B cells in addition to gut plasma cells, which had either a short-lived or intermediate-lived phenotype (Figure 5C). These clonally related cells in blood and gut likely result from ongoing immune reactions accompanied by continued BCR diversification in patients with untreated CeD.

**Figure 5.**
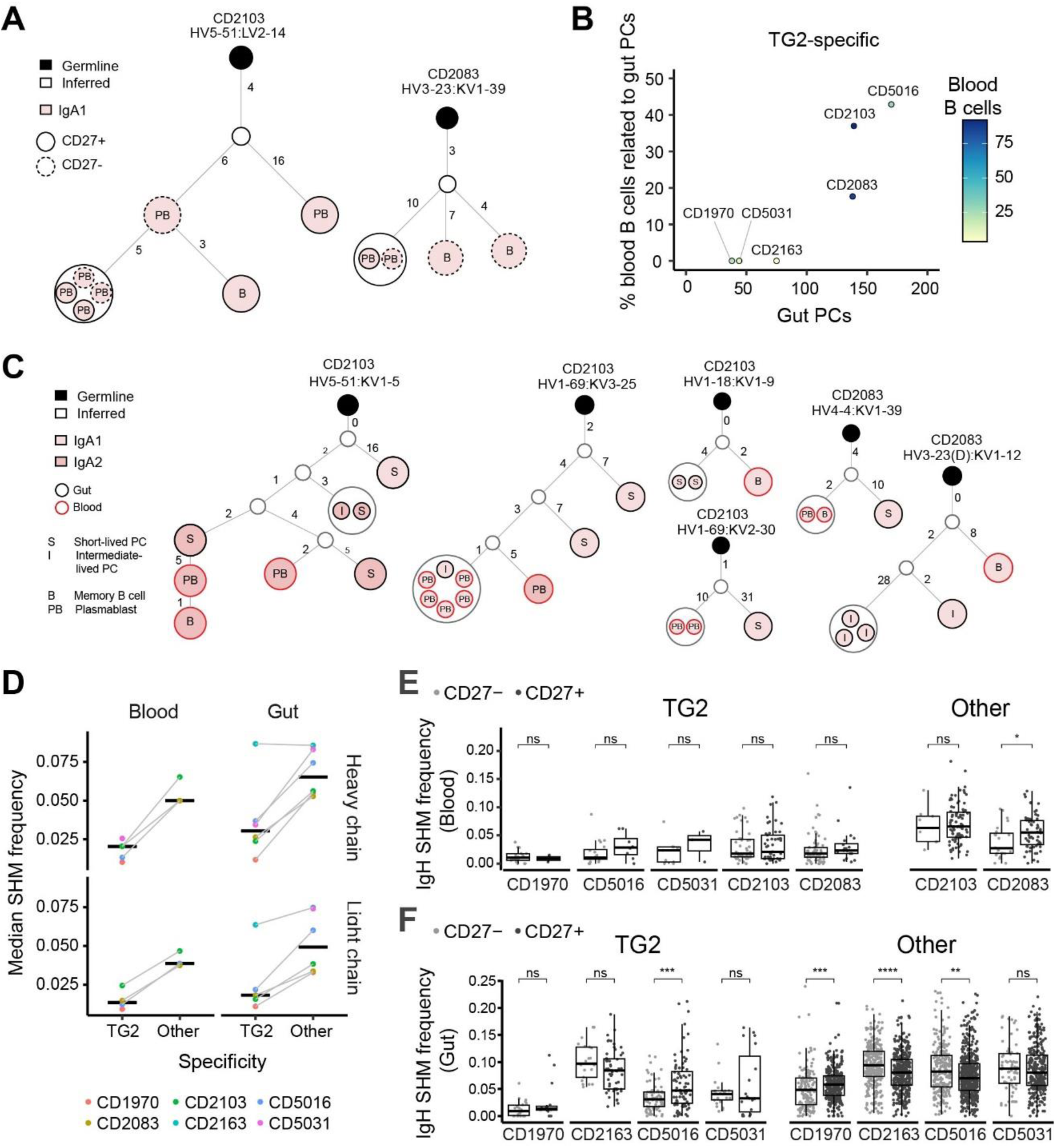
Clonal relationships and Ig mutations in circulating B cells and gut plasma cells. (A) Examples of lineage trees showing clonal relationships between circulating TG2-specific memory B cells (MBC) and plasmablasts (PB) with or without surface CD27. (B and C) Quantification of overlap (B) and examples of lineage trees (C) showing clonal relationships between TG2-specific gut plasma cells (PCs) and IgA+ B cells in blood of six untreated CeD patients. PCs were defined as either short-lived (CD19+CD45+) or intermediate-lived (CD19-CD45+) based on flow cytometry staining. (D) Frequency of heavy and light chain variable region mutations introduced by somatic hypermutation (SHM) among TG2-specific and non-TG2-specific (other) IgA+ blood B cells or gut plasma cells of six untreated CeD patients. (E and F) Comparison of mutation levels between CD27+ and CD27- IgA cells in blood (E) and gut biopsies (F). Centers indicate medians, and statistical significance was evaluated by a Mann-Whitney U test. *p < 0.05.

As previously reported, TG2-specific tissue-resident plasma cells harbored fewer Ig mutations than other gut plasma cells.^2,22^ We here show that the difference between TG2-specific and non-TG2- specific cells also holds true for circulating IgA cells (Figure 5D). The TG2-specific plasmablasts had particularly few mutations, suggesting that these cells represent an early differentiation stage following activation of naïve B cells (Figure S4D). Although we observed substantial variation between individual cells, the CD27- cells in blood tended to have fewer Ig mutations than the CD27+ ones (Figure 5E). This difference was observed for both TG2-specific and other IgA-switched cells in accordance with previous reports on circulating DN cells in healthy individuals.^13,23–25^ Curiously, we did not see a similar difference between CD27+ and CD27- cells among gut plasma cells (Figure 5F). The explanation for this apparent discrepancy is probably that the CD27- population is enriched for CD19+ short-lived cells, which have been shown to harbor more mutations than the more long-lived populations.^3^

### Circulating TG2-specific B cells are activated and gut-homing

Transcriptomic profiling of circulating IgA-switched memory B cells and plasmablasts confirmed that TG2-specific cells in untreated CeD have lower expression of CD27 than non-TG2-specific cells (Figures 6A and 6B). Further, the TG2-specific cells showed expression of homing molecules associated with migration toward inflamed sites (CXCR3) and the mucosa of the small intestine (integrin α4β7 and CCR9). Conversely, they had lower expression of homing molecules controlling migration to lymphoid tissues (CXCR4 and CD62L). Compared with other memory B cells, the TG2-specific cells had higher expression of J-chain – a sign of activation and plasma cell differentiation.^26^ They also expressed other activation markers such as CD38, CD86 and CD95 as well as CD11c, which is found on some activated B-cell subsets.^27–30^ Thus, the TG2-specific cells in the memory B-cell cluster appear to be activated rather than resting, and they likely represent an early stage of plasma cell differentiation (Figure 6B).

**Figure 6.**
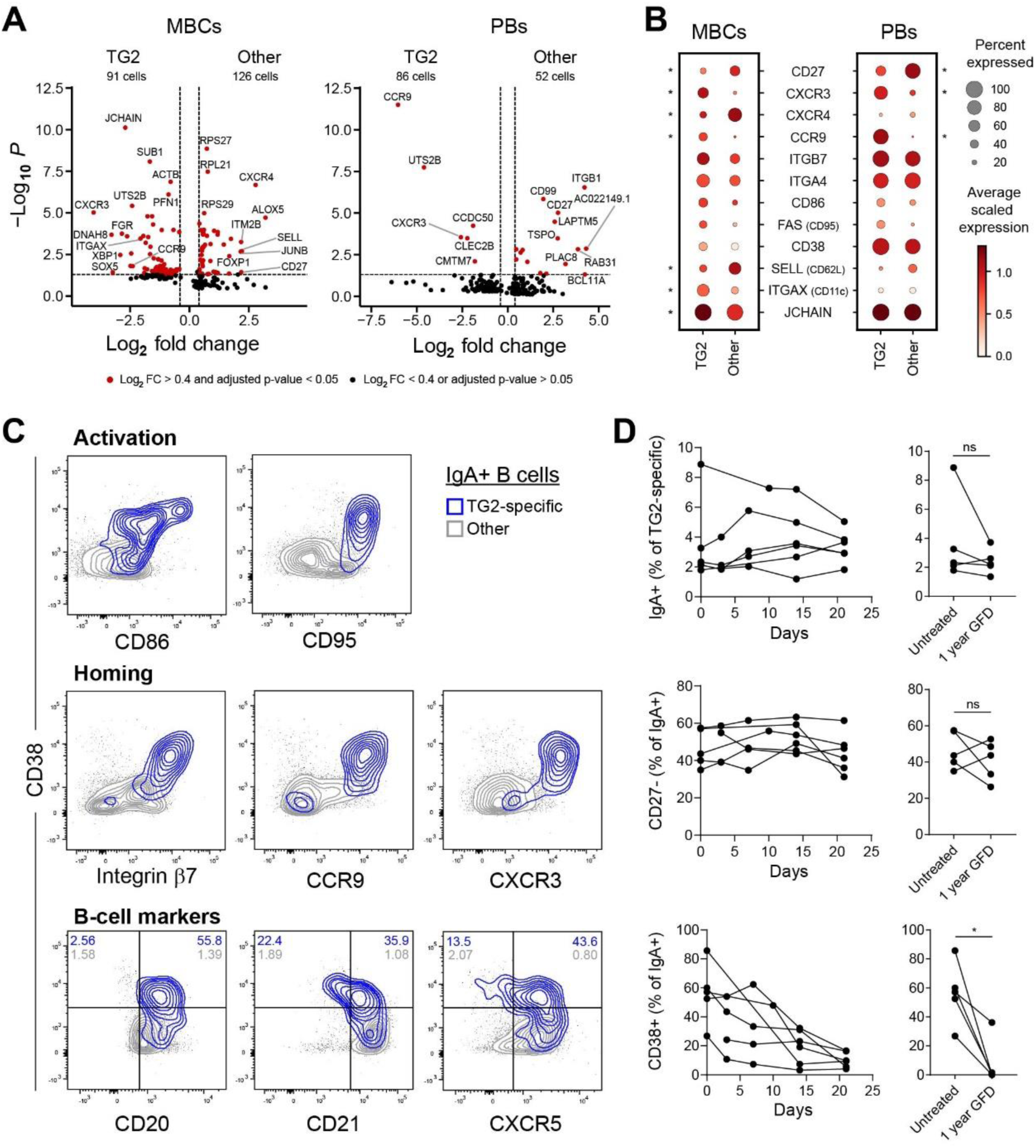
Gene expression profile of circulating TG2-specific B cells. (A) Volcano plots showing identification of differentially expressed genes between TG2-specific and non-TG2-specific (other) IgA+ B cells belonging to either the memory B cell (MBC) or plasmablast (PB) cluster (Figure 4E). (B) Expression of selected genes associated with migration and activation in the indicated cell populations. Genes with statistically significant (p < 0.05) differential expression are indicated with *. (C) Representative flow cytometry plots of circulating IgA+ B cells in an untreated CeD patient showing relationship between surface CD38 and markers associated with activation, homing or MBCs. (D) Change in number and phenotype of TG2-specific IgA+ B cells in six patients during the first three weeks or after one year of gluten-free diet (GFD) as determined by flow cytometry. Difference between time points was evaluated by a paired t test. *p < 0.05.

To confirm that most TG2-specific B cells have an activated, gut-homing phenotype, we characterized B-cell surface marker expression in patients with active CeD by flow cytometry (Figures 6C and S4). Again, we observed that many of the TG2-specific IgA cells expressed CD38 in combination with homing and activation markers, which indicate that the cells are migrating to the inflamed gut and are destined to become tissue-resident plasma cells. Not all CD38+ cells were bona fide plasmablasts, as some of the cells retained expression of B-cell markers such as CD20, CD21 and CXCR5 that are typically lost upon differentiation (Figure 6C). In agreement with our scRNA-seq data, most of the TG2- specific cells had either a plasmablast (CD20-CD21-CD38^Hi^CXCR5-) or an activated memory B-cell (CD20+CD21+CD38+CXCR5+) phenotype. In both cases, the cells were largely negative for CD27, but they did not fall into the category of DN2 cells, which is characterized by high expression of CD19 and CD11c in combination with low expression of CD21 and CXCR5 (Figure S5).^31–34^

We next addressed the fate of circulating autoreactive B cells in CeD by following TG2-specific IgA cells in patients from they were diagnosed and for a period after they started on a gluten-free diet (Figures 6D and S6). The cells did not disappear within the first few weeks of gluten-free diet. In fact, some patients showed an initial increase in the frequency of TG2-specific IgA cells followed by a gradual decline. The appearance of additional cells in the blood during the first few weeks after removal of the driving antigen (gluten) might be explained by contraction of lymphoid tissues harboring TG2- specific memory B cells.^35^ The frequency of CD27- cells did not appear to change during the first phase of gluten-free diet. However, the frequency of CD38+ TG2-specific cells rapidly dropped (Figure 6D). Since CD38 expression was correlated with high levels of integrin β7, the disappearance of CD38+ cells from blood is likely due to their migration into the gut mucosa in combination with a stop in generation of new activated cells (Figure S6). Collectively, our results indicate that circulating TG2-specific B cells with an activated phenotype are continuously generated during active CeD and that these cells are migrating to the gut to become tissue-resident plasma cells.

### TG2-specific IgA cells in untreated CeD are generated from naïve B cells

To gain insight into the origin of autoreactive B cells in CeD, we aimed to recreate the phenotype of TG2-specific IgA cells in vitro by using our B-cell culture system to mimic T cell-dependent activation. Culturing of naïve B cells with transfected fibroblasts led to efficient activation accompanied by downregulation of surface IgD (Figure 7A). By supplementing the cultures with retinoic acid, a substantial number of IgA-switched cells could be obtained from naïve B cells (Figures 7B and S7). When comparing such naïve-derived IgA cells with cultures of isolated IgA cells that were either positive or negative for CD27 to begin with, we observed that only the naïve cells gave rise to a substantial population of cells that were negative for CD27 and positive for CD38 and thus phenocopied the TG2-specific IgA cells seen in CeD (Figure 7C). These results suggest that most of the autoreactive B cells we observe in active CeD are generated through activation of naïve B cells rather than stimulation of resting memory cells.

**Figure 7.**
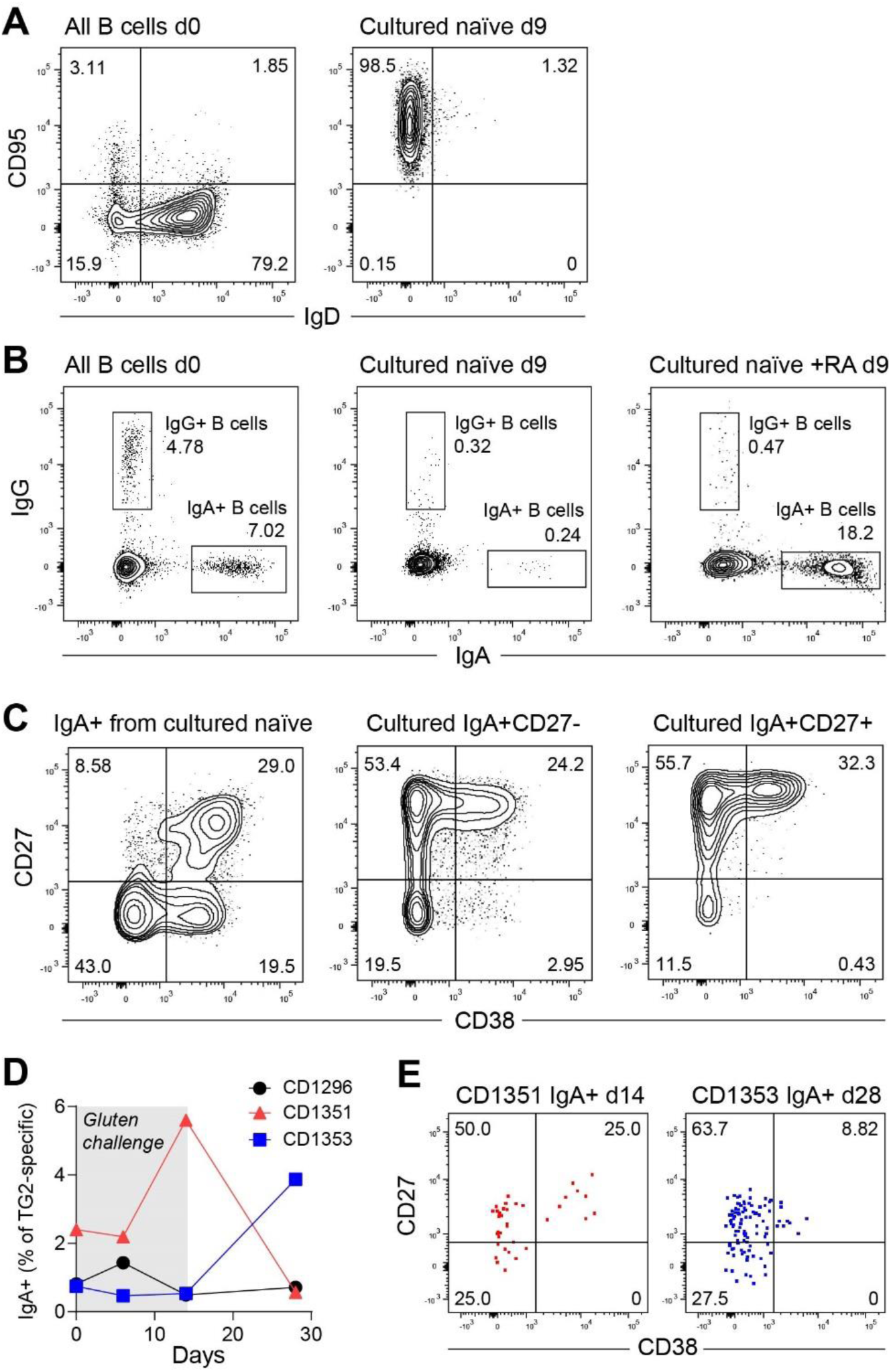
Phenotype of B cells activated in vitro or in vivo. (A-C) Representative flow cytometry plots showing staining of B cells from peripheral blood of a healthy donor before (d0) or after (d9) nine days of culture with transfected fibroblasts expressing human BAFF, CD40L and IL-21. The cultured B cells were analyzed for activation (A), class-switching (B) and similarity to TG2-specific IgA+ B cells in CeD (C). To induce IgA-switching, cultures of naïve B cells were supplemented with retinoic acid (RA). (D and E) Induction of a TG2-specific B-cell response in treated patients undergoing a 14-days oral gluten challenge. The frequency of IgA-switched TG2-specific B cells in peripheral blood was followed during and after the challenge by flow cytometry (D). In the two patients, who responded to the challenge, expression of surface CD27 and CD38 by the TG2-specifc cells was assessed at the peak of the response (E).

We next aimed to address the initial stages of B-cell activation in vivo by following autoreactive B cells in treated CeD patients, who were challenged with a gluten-containing diet.^36^ In two out of three patients, we observed an increase in the number of TG2-specific IgA cells – in one patient on day 14 and in the other on day 28 after initiation of the challenge (Figure 7D). The rather slow B-cell response is consistent with a previous report showing that it takes several weeks to generate TG2-specific serum antibodies upon gluten challenge of treated CeD patients,^37^ possibly because the response depends on induction of mucosal damage and release of intracellular TG2.^38,39^ In both responding patients, a small population of CD38+ cells could be detected, indicating B-cell activation (Figure 7E). However, these cells were positive for CD27 and, hence, do not reflect the population of activated CD27- cells observed in untreated CeD. These results suggest that the acute response during gluten challenge is different from the chronic response of untreated CeD. It is likely that the acute response primarily recruits resting memory B cells, whereas the chronic response relies on activation of naïve B cells for continuous plasma cell generation.

## DISCUSSION

A hallmark of CeD is massive accumulation of small intestinal plasma cells that secrete TG2-specific autoantibodies.^40^ We here describe the corresponding population of circulating TG2-specific IgA B cells in blood samples of CeD patients. These cells were present in patients with untreated CeD but not in healthy donors, and their frequency was greatly reduced in most patients on a long-term gluten-free diet. The TG2-specific B cells in blood showed a strong clonal relationship with TG2-specific gut plasma cells, indicating that they share a common induction site. Notably, both populations carried relatively few Ig mutations compared with other IgA-switched cells.

In untreated CeD, most circulating TG2-specific B cells were either plasmablasts or activated memory B-cells. Both phenotypes were characterized by expression of CD38, but they differed in expression of typical B-cell markers such as CD20, CD21 and CXCR5. In addition, the cells expressed homing markers consistent with migration to the inflamed small intestine. In agreement with a previous study, circulating TG2-specific IgA cells included classical CD27+ memory B cells.^41^ The majority of the TG2- specific cells, however, were negative for CD27, making them different from most other IgA-switched cells. The lack of CD27 expression was observed for both B cells/plasmablasts in blood and plasma cells in duodenal biopsies of untreated CeD patients. Despite being negative for CD27, the phenotype of TG2-specific cells did not match DN2 cells, which are a population of activated CD27- B cells found to be elevated during chronic infections and in some autoimmune diseases, most notably in systemic lupus erythematosus (SLE).^33^ DN2 cells are also referred to as tissue-like B cells^31,42^ or atypical B cells^15^ in humans and age-associated B cells in mice.^43^ They are believed to be poised for plasma cell differentiation, but unlike the TG2-specific cells we describe here, DN2 cells do not express the plasma cell marker CD38.

Rather than belonging to the DN2 subset, activated TG2-specific memory B cells in CeD show similarity to another DN subset – the DN1 cells. Unlike DN2 cells, DN1 cells are positive for CXCR5 and CD21.^33^ In healthy individuals, circulating DN1 cells were shown to be enriched for IgA-switched B cells, and they express many of the same markers that we find in the TG2-specific memory cells, including J- chain and CD38.^30,44^ Based on their transcriptomic profile and similarity to classical memory B cells, DN1 cells were proposed to represent precursors of class-switched memory cells that have not yet acquired CD27 expression.^33^ However, expression of typical plasmablast/plasma cell genes, including *JCHAIN*, *CD38* and *XBP1*, suggests that circulating TG2-specific memory B cells are activated plasmablast/plasma cell precursors. Further, we and others have shown that fully differentiated plasmablasts and plasma cells can be negative for CD27.^44^ Thus, CD27- cells are not simply precursors of CD27+ effector cells.

In SLE, the frequency of circulating DN2 cells is greatly increased compared to healthy individuals, and excessive DN2 formation depends on B-cell stimulation via TLR7 in addition to T cell-derived signals.^33,45^ The DN2 population was shown to harbor autoreactive B cells, although autoreactivity was not restricted to this subset.^33,34^ Further, DN2 cells have also been described in the context of acute infection and vaccination, and their formation could thus represent a normal part of ongoing immune reactions that also involve classical class-switched B cells expressing CD27.^46–48^ Targeted characterization of antigen-specific B cells in other autoimmune diseases indicates that circulating autoreactive B cells typically express CD27 and thus seem to be generated through classical memory B-cell responses.^49,50^ Conversely, we here demonstrate that most TG2-specific IgA cells in untreated CeD lack CD27 expression.

Activation of TG2-specific B cells and formation of TG2-specific autoantibodies depend on B-cell interactions with gluten-specific CD4+ T cells. Circulating and gut-resident gluten-specific CD4+ T cells were shown to have a T peripheral helper (Tph)-like phenotype characterized by expression of PD-1 and the B-cell helper cytokines IL-21 and CXCL-13 in combination with a lack of CXCR5.^51^ Such T cells are believed to be involved in T cell-B cell interactions outside of GCs.^52^ A primarily extrafollicular activation route is supported by the low mutation levels in TG2-specific B cells and the limited number of long-lived plasma cells in CeD patients on a long-term gluten-free diet.^3^ Thus, based on the phenotypes of both B cells and T cells, we propose that TG2-specific B cells in CeD are generated through T cell-dependent activation at extrafollicular sites. Similarly, circulating DN2 cells in SLE were proposed to be derived from extrafollicular activation of naïve B cells^33^ in agreement with a non-GC phenotype of DN2 cells in human tonsils^28^ and prominent extrafollicular B-cell responses in mouse models of SLE.^53,54^ All DN cells might therefore originate from extrafollicular B-cell activation, where the choice between DN1 and DN2 fates could be controlled by TLR7 ligation. Hence, CD27 is a potential marker of GC-derived cells in humans.^55^ Importantly, though, we found CD27+ and CD27- cells with identical BCR sequences, suggesting that CD27 expression on activated B cells could be subject to dynamic changes.

The fate of activated B cells is influenced by the availability of antigen.^56^ For example, immunization can drive prolonged GC reactions through accumulation and persistence of antigen on follicular dendritic cells.^57^ In CeD, we have proposed that shed enterocytes are the source of self-antigen through release of intracellular TG2 into the gut lumen followed by uptake in gut-associated lymphoid tissues (GALT) such as Peyer’s patches.^58^ TG2 that enters Peyer’s patches via microfold cells or conduits^59^ will likely accumulate in the subepithelial dome (SED) due to strong interactions between TG2 and components of the extracellular matrix.^60^ Thus, TG2 that is taken up from the gut lumen will not reach the underlying B-cell follicle and be able to drive GC reactions. However, the SED contains both B cells and CD4+ T cells, and it is a major site for IgA class switching.^61^ Hence, it is possible that interactions between TG2-specific B cells and gluten-specific T cells take place in the SED and that the distribution of TG2 within GALT is thereby responsible for skewing of B-cell activation toward an extrafollicular rather than GC-dependent response.

In summary, we have shown that circulating TG2-specific IgA+ B cells in CeD are the precursors of autoantibody-secreting gut plasma cells. The cells were negative for CD27, and they had a plasmablast or activated DN1-like phenotype. Due to their low mutation levels and resemblance to naïve B cells, we propose that the autoreactive IgA cells are generated through extrafollicular interactions involving TG2-specific naïve B cells and gluten-specific CD4+ T cells during chronic exposure to dietary gluten. The circulating IgA-switched cells that we describe here are likely activated in GALT and travel via the blood to seed the small intestinal lamina propria as tissue-resident plasma cells. By employing an antigen-specific approach, our study provides insight into the function and origin of the enigmatic DN population of human B cells and the mechanisms that drive autoantibody formation in CeD.

## Limitations

Based on the phenotype of circulating TG2-specific IgA cells, we propose that autoreactive B cells in active CeD are derived from recruitment of naïve B cells to an extrafollicular, T-cell dependent immune response. To obtain direct evidence for this model, it would be necessary to follow the TG2-specific repertoire in individuals with chronic CeD and show that new clones replace old ones over time. In addition, to establish that B cells undergo extrafollicular activation, it would be necessary to obtain biological samples from inductive sites (presumably GALT) and assess the location of antigen-specific B cells and T cells in situ. The in vivo availability of antigen (TG2 and gluten), including possible accumulation of extracellular TG2 in the SED, should also be addressed in order to obtain further mechanistic insight into the gluten-dependent formation of autoantibodies in CeD.

## ACKNOWLEDGEMENTS

We are thankful to all patients who have donated biological material to this study and to the staff at the Endoscopy Units at Oslo University Hospital and Akershus University Hospital for help with sample collection. Flow cytometry and cell sorting experiments were conducted at the Flow Cytometry Core Facility, Oslo University Hospital, and next generation sequencing was performed at the Norwegian Sequencing Centre, University of Oslo. Raw sequencing data were stored and processed on the TSD (Tjeneste for Sensitive Data) facilities, owned by the University of Oslo. This work was supported by grants from the South-Eastern Norway Regional Health Authority (projects 2016113 and 2020027), Stiftelsen KG Jebsen (project SKGJ-MED-017) and the University of Oslo World-leading research program on human immunology (WL-IMMUNOLOGY).

## AUTHOR CONTRIBUTIONS

Conceptualization, R.I. and L.M.S.; Methodology, R.I., I.L., A.C., and L.F.R.; Investigation, R.I., I.L., L.S.H., and A.C.; Resources, A.C., L.F.R., J.J., and K.E.A.L.; Writing – Original Draft, R.I., and I.L.; Writing – Review & Editing, all authors; Supervision, L.M.S.; Funding Acquisition, L.M.S.

## MATERIALS AND METHODS

### Human samples

Duodenal biopsies and/or peripheral blood were collected from adult CeD patients (n=50) with a median age of 32 years (range 18-80). Fifty nine percent were females, and the patients were either untreated or had been on a gluten-free diet for various periods of time. The diagnosis of CeD was established based on the guidelines of the European Society for the Study of Coeliac Disease,^62^ and mucosal damage was assessed by Marsh score classification.^63^ In one set of experiments, samples were obtained from three treated patients who had been challenged with 5.7 g of gluten per day for 14 days.^36^ Biopsy samples from two control donors were obtained from individuals undergoing endoscopy but where CeD was excluded. Blood samples from healthy donors were obtained as buffy coats from Oslo Blood Bank. All participants gave their informed consent. Approval of experimental procedures was obtained from the Regional Committee for Medical and Health Research Ethics of South-Eastern Norway (REK ID 6544 and 13742).

### Recombinant proteins

Human TG2 used for flow cytometry contained an N-terminal His6-tag followed by an Avi-tag and was expressed in *Eschericia coli* BL21 (DE3) as described.^64^ The control antigen GST containing the same tags as TG2 was generated by PCR, using the pGEX-3X vector as template followed by subcloning into the pET-28a expression vector between the NcoI and HindIII restriction sites. Both proteins were purified from *E. coli* lysates by Ni-NTA affinity chromatography followed by site-specific biotinylation using BirA biotin-protein ligase (Avidity). The biotinylated proteins were further purified by anion-exchange chromatography using a Mono Q column (GE Healthcare) and stored in 20 mM Tris-HCl, pH 7.3, 300 mM NaCl, 1 mM DTT, 1 mM EDTA. TG2 used for coating in ELISA contained a C-terminal His6-tag and was expressed in Sf+ insect cells as previously described^60^.

### Flow cytometry

Duodenal biopsies were collected in ice-cold RPMI-1640 and processed into single-cell suspensions by treatment with 2 mM EDTA in 2% (v/v) FBS/PBS followed by digestion with collagenase (Sigma) and cryopreservation as previously described.^22^ PBMCs were isolated from peripheral blood by density gradient centrifugation followed by cryopreservation. To identify antigen-specific B cells and plasma cells, duodenal single-cell suspensions or PBMCs were stained with tetramerized antigens that were generated before each experiment as follows: biotinylated TG2 and biotinylated GST were incubated with streptavidin-PE (Thermo) and streptavidin-PE/Cy7 (Thermo), respectively, using a 1.5-fold molar excess of biotinylated molecules over biotin binding sites in 2% FBS/PBS. After incubation for 1 h, the antigen-streptavidin complexes were mixed together and added to the cells to reach a final antigen concentration of 1-3 µg/ml followed by incubation on ice. In one set of experiments, staining for endogenous BCR-bound TG2 was performed prior to staining with recombinant TG2. Antibodies specific for individual surface markers were either added directly or after antigen-specific enrichment. To enrich for antigen-binding cells, labeled cell suspensions were incubated with anti-PE microbeads (Miltenyi) on ice. The cells were then washed and passed over magnetized MS or LS Columns (Miltenyi) followed by washing and elution of captured cells in 2% FBS/PBS. Enriched and non-enriched fractions were analyzed on an LSRFortessa (BD) or a FACSymphony A5 (BD) instrument. Dead cells were identified by labeling with LIVE/DEAD fixable Aqua (Thermo) prior to incubation with antigens and antibodies.

### Antibodies used for flow cytometry

Anti-human CCR9-BUV395 (clone C9MAB-1, BD), anti-human CD3-BV510 (clone OKT3, BioLegend), anti-human CD3-BV605 (clone UCHT1, BioLegend), anti-human CD14-BV510 (clone M5E2, BioLegend), anti-human CD14-BV605 (clone M5E2, BioLegend), anti-human CD11c-BUV805 (clone B-ly6, BD), anti-human CD11c-BV605 (clone 3.9, Biolegend), anti-human CD19-Pacific blue (clone HIB19, BioLegend), anti-human CD19-PerCP/Cy5.5 (clone HIB19, BioLegend), anti-human CD19-APC/Cy7 (clone HIB19, BioLegend), anti-human CD20-BUV496 (clone 2H7, BD), anti-human CD20-BV605 (clone 2H7, BioLegend), anti-human CD21-BV421 (clone 1048, BD), anti-human CD21-APC (clone Bu32, BioLegend), anti-human CD24-BV605 (clone ML5, BioLegend), anti-human CD27-BV421 (clone O323, BioLegend), anti-human CD27-BV711 (clone M-T271, BioLegend), anti-human CD27-PE/Cy7 (clone LG.7F9, eBioscience), anti-human CD27-APC (clone M-T271, BioLegend), anti-human CD27-APC/Cy7 (clone O323, BioLegend), anti-human CD38-BUV563 (clone 2B7, BD), anti-human CD38-BV605 (clone HIT2, BioLegend), anti-human CD38-FITC (clone HB7, eBioscience), anti-human CD38-PerCP-Cy5.5 (clone HIT2, Biolegend), anti-human CD62L-R718 (clone DREG-56, BD), anti-human CD86-BV785 (clone IT2.2, BioLegend), anti-human CD95-BUV737 (clone DX2, BD), anti-human CD95-Alexa Fluor 700 (clone DX2, BioLegend), anti-human CXCR3-PE-CF594 (clone 1C6/CXCR3, BD), anti-human CXCR4-BV605 (clone 12G5, BD), anti-human CXCR5-PE/Cy5 (clone J252D4, BioLegend), anti-human CXCR5-Alexa Fluor 647 (clone J252D4, BioLegend), anti-human IgA-FITC (clone IS11-8E10, Miltenyi), goat anti-human IgA-PE (Southern Biotech), anti-human IgA-APC (clone IS11-8E10, Miltenyi), anti-human IgA-APC-Vio770 (clone IS11-8E10, Miltenyi), goat anti-human IgD-FITC (Southern Biotech), anti-human IgD-PerCP/Cy5.5 (clone IA6-2, BioLegend), anti-human IgD-Alexa Fluor 700 (clone IA6-2, BioLegend), anti-human IgG-APC (clone M1310G05, BioLegend), anti-human IgM-BV510 (clone MHM-88, Biolegend), goat anti-human IgM-FITC (Southern Biotech), anti-human Integrin β7-BV650 (clone FIB504, BD), anti-human Integrin β7-APC (clone FIB504, BioLegend), rabbit anti-human TG2 (Pacific Immunology), goat anti-rabbit IgG-Alexa488 (Thermo Fisher), anti-biotin-PE (clone 1D4-C5, Biolegend).

### ELISAs

To assess production of TG2-specific and total antibodies in B-cell cultures, 3 µg/ml recombinant human TG2 or 2 µg/ml goat anti-human Ig (Southern Biotech) was coated in PBS followed by incubation with culture supernatants diluted in PBS supplemented with 0.1% (v/v) Tween-20 (PBST) and 1% (w/v) BSA. Bound antibodies were subsequently detected with alkaline phosphatase-conjugated goat anti-human isotype-specific secondary antibodies. Total antibody concentrations were determined based on standard curves generated with recombinant anti-TG2 monoclonal antibodies (clone 679-14-E06) expressed with the relevant heavy chain constant regions.^65^

To detect secretion of IL-21 from transfected fibroblasts, mouse anti-human IL-21 (clone 3A3-N2, eBioscience) was coated at 2 µg/ml in PBS followed by incubation with cell supernatants diluted in 1% BSA/PBST. Bound IL-21 was subsequently detected with mouse anti-human IL-21-biotin (clone 2B2-G20, eBioscience) followed by alkaline phosphatase-conjugated streptavidin (Southern Biotech). The signals were compared to a standard curve generated with recombinant human IL-21 (R&D Systems).

### ELISPOT

Spontaneous antibody secretion from IgA-switched B cell populations was assessed in MultiScreen 96-well filter plates (Millipore) that were coated with 2 µg/ml goat anti-human Ig (Southern Biotech) in PBS. The wells were blocked with 1% BSA/PBS before addition of IgA B cells (CD3^-^ CD14^-^CD19^+^IgD^-^IgA^+^CD27^+/-^CD38^+/-^) that had been bulk-sorted from stained PBMCs using a FACSAriaIII instrument (BD). The cells were incubated overnight at 37°C, 5% CO_2_, in B-cell medium (10% FBS/RPMI-1640 supplemented with 10 mM HEPES, pH 7.4, 1 mM sodium pyruvate, non-essential amino acids and penicillin-streptomycin) followed by extensive washing with PBS and PBST. Bound antibodies were subsequently detected by incubation with alkaline phopsphatase-conjugated goat anti-human IgA (Sigma) in 1% BSA/PBST followed by washing and addition of BCIP/NBT substrate (Bio-Rad). Pictures of individual wells were captured using an ImmunoSpot analyzer (CTL).

### Generation of transfected fibroblasts

To establish a feeder cell line for culturing of single B cells, mouse NIH/3T3 fibroblasts were transfected with NotI-linearized pUNO1 vectors (Invivogen) encoding human BAFF (CD257) and human CD40L (CD154) by electroporation of 10^7^ cells with 10 µg of each plasmid, using an ECM 830 electroporation system (BTX). Two days later, the transfected cells were transferred to 10% FBS/DMEM containing 5 µg/ml blasticidin. To check for expression of the transgenes, cells that had been detached with 0.25% (w/v) trypsin-EDTA (Thermo) were allowed to recover for 2 h at 37°C, before surface expression was assessed with anti-human CD257-PE (clone 1D6, BioLegend) and anti-human CD154-APC (clone 24-31, BioLegend). Double positive cells were bulk-sorted using a FACSAriaIII instrument (BD) and further cultured. To introduce expression of IL-21, the human cDNA sequence (Invivogen) was subcloned into the pcDNA3 vector between the HindIII and EcoRV restriction sites followed by transfection of 5 µg plasmid into BAFF/CD40L double positive fibroblasts using Lipofectin reagent (Thermo). Two days later, the cells were transferred to medium containing 250 µg/ml geneticin (Thermo) and further cultured. Single clones were then established by limiting dilution. After 11 days, secretion of IL-21 into the culture medium was assessed by ELISA. One clone (2B09) that was positive for IL-21 and supported B-cell activation was selected for further testing and use in B-cell culture experiments. Expression of all three transgenes was found to be stable after culturing the 2B09 cells for several weeks in the absence of selective antibiotics.

### B-cell cultures

The day before initiation of an experiment (day -1), triple-transfected 2B09 cells or untransfected NIH/3T3 fibroblasts were irradiated (25 Gy) and plated at 5,000 cells per well in 96-well flat-bottom plates in 100 µl B-cell medium. In initial experiments, total B cells were isolated from PBMCs of a healthy donor by magnetic depletion of non-B cells using B-cell Isolation Kit II (Miltenyi). Before addition of the B cells, the fibroblast medium was replaced with 100 µl fresh B-cell medium. The B cells were then added to the fibroblasts at 0.5 cells per well and cultured at 37°C, 5% CO_2_. For culturing of TG2-binding or non-TG2-binding IgA cells, single cells were sorted from patient PBMCs directly into 2B09-coated wells using a FACSAriaIII instrument (BD). Part of the medium was replaced with 100 µl fresh B-cell medium three times during the culture period: 50 µl was replaced on day 2 or 3, and 80 µl was replaced on day 4-6 and again on day 7 or 8. On day 9 or 10, the cell supernatant was collected for assessment of antibody secretion by ELISA.

For phenotypic characterization of cultured B cells, 2B09 fibroblasts were plated in 24-well plates at 10,000 cells per well in 1 ml B-cell medium without irradiating the cells. The next day (day 0), different B cell populations were bulk sorted from PBMCs of healthy donors using a FACSMelody instrument (BD). The fibroblast medium was replaced with 1 ml fresh B-cell medium before addition of 4000 naïve (CD3^-^CD14^-^CD19^+^IgD^+^CD27^-^) or 500 IgA-switched (CD3^-^CD14^-^CD19^+^IgD^-^IgA^+^CD27^+/-^) B cells in 500 µl B-cell medium. To induce IgA-switching, some of the naïve B-cell cultures were supplemented with retinoic acid (Sigma) to a final concentration of 100 nM. On day 2 or 3, a new layer of feeder cells was prepared from 10,000 2B09 fibroblasts in 1 ml B-cell medium. The cultured B cells were transferred to the new layer by first removing 800 µl of the supernatant and resuspending the cells in the remaining medium. This procedure was repeated on days 5 and 7 or on day 6. On day 9, the cells were collected and analyzed by flow cytometry.

### Ig sequencing

After screening of single-B-cell cultures by ELISA, positive wells were selected, and the cells were collected by resuspension in 10 mM EDTA/PBS. Obtained cell pellets were snap-frozen on dry ice and stored at -80°C until further analysis. After thawing on ice, the cells were resuspended in TCL lysis buffer (Qiagen) supplemented with 1% (v/v) 2-mercaptoethanol, and total RNA was purified by use of RNAClean XP beads (Beckman Coulter). Heavy and light chain variable regions were then amplified using a nested RT-PCR approach with previously reported primers for specific amplification of IgA antibody sequences.^66,67^ The PCR products were purified and Sanger-sequenced using constant region-specific reverse primers.

### Droplet-based scRNA-seq

TG2-binding IgA plasma cells were identified by flow cytometry analysis of duodenal biopsy single-cell suspensions as described above with the exception that TG2 tetramers were generated with DNA-barcoded streptavidin-PE (TotalSeq-C0951, BioLegend). In addition, a barcoded anti-human CD27 antibody (TotalSeq-C0154, BioLegend) was included together with fluorophore-conjugated antibodies. The cells were incubated with FcR Blocking Reagent (Miltenyi) prior to addition of antibodies and antigens. Following incubation on ice, the cells were washed, and TG2-binding and non-TG2-binding IgA plasma cells were bulk-sorted into empty microtubes that had been coated with 1% BSA/PBS using a FACSAriaIII instrument (BD). For each of four untreated CeD patients, TG2-binding and non-TG2-binding cells were combined. In addition, two of the samples contained 500-1000 hash-tagged cells that were sorted for an unrelated study. In the end, barcoded cDNA from a total of 5,000-10,000 cells per sample was generated using the 10x Genomics Chromium Controller. Single cell 5’ gene expression, V(D)J-enriched and cell surface protein libraries were generated using Chromium single cell kits (v1.0) according to the instructions from the manufacturer (10x Genomics). The libraries were pooled prior to sequencing on a NovaSeq 6000 instrument (Illumina) using the following configuration: read 1: 26 cycles, read 2: 89 cycles, index read 1: 8 cycles.

### Smart-seq2

TG2-binding and non-TG2-binding IgA B cells were identified by flow cytometry analysis of freshly isolated PBMCs as described above. Single cells of two untreated CeD patients were sorted into 96-well plates containing 2 µl of scRNA-seq catch buffer (0.2% vol/vol Triton X-100 [Sigma] in H_2_O with 2 U/μl RNase inhibitor [New England Biolabs]) per well using a FACSAriaIII instrument (BD). The cells were then snap-frozen on dry ice and stored at -80°C until library preparation. Reverse transcription and cDNA pre-amplification were performed using a modified version of Smart-seq2^20^ with 0.5 μl/well SMARTScribe Reverse Transcriptase (Clontech) for reverse transcription and 24 PCR cycles for cDNA pre-amplification. Pre-amplified cDNA was purified with 20 μl Ampure XP beads (Agencourt) per well, tagmented using an inhouse-produced Tn5 transposase^68^ and dual indexed with Nextera (XT) N7xx and S5xx index primers (final concentration 125 nM). Four plates were pooled together and sequenced on a NextSeq500 instrument (Illumina) with 75 bp paired-end reads in high-output mode.

### scRNA-seq quality control

*Smart-seq2.* Raw reads were trimmed for adapter- and low-quality sequences with Cutadapt v.1.18^69^ through Trim Galore v0.6.1 in paired-end mode. Transcript expression was quantified with Salmon v0.11.3^70^ using cDNA sequences from GRCh38.94 and k-mer length 25 and aggregated to gene level. We prepared a transcript-length-corrected gene expression matrix using tximport v1.8.0.^71^ The cells were subjected to quality control based on the following criteria: 1500-7500 detected genes, 100000-3000000 reads, < 8% mitochondrial genes, > 50% reads mapping to the reference transcriptome and a productively rearranged BCR heavy chain of the IgA isotype reconstructed by BraCeR (see **BCR reconstruction and clonal assignments**). Quality control was performed in R version 3.5.3 using the scater package.^72^ In total 355 cells remained after quality control.

*Droplet-based scRNA-seq.* The gene expression, V(D)J and cell surface protein data were processed through Cell Ranger 6.0.2 with the *multi* and *aggr* functions using the pre-built Cell Ranger references GRCh38 version 2020-A for gene expression and GRCh38-alts-ensembl-5.0.0 for V(D)J analysis. Quality control of the output of Cell Ranger was performed using Seurat v4.3.0.1^73^ in R 4.3.1 based on the following criteria: 200-2000 detected genes, 2000-25000 Unique Molecular Identifiers (UMIs), < 10% mitochondrial genes, > 10% Ig genes and a productively rearranged BCR heavy chain of the IgA isotype reported in the “airr_rearrangement.tsv” files generated by Cell Ranger *multi* for each patient. Irrelevant cells (see above) were identified and removed based on hash-tag labeling, which was normalized by centered log-ratio transformation. In total 3146 cells passed quality control.

### scRNA-seq data analysis

*Smart-seq2.* The smart-seq2 gene expression analysis was performed using scanpy v1.9.1^74^ in python 3.7 unless otherwise stated. All Ig genes were discarded from the transcriptional analysis before normalization to avoid masking of non-Ig-related transcriptional differences. Genes detected in more than 5 cells were retained. The gene expression matrix was normalized by total counts per cell and logarithmized as *X* = ln(*X* + 1). Highly variable genes were identified with the *highly_variable_genes* function (min_mean=0.1, max_mean=10, min_disp=0.25). Unwanted variation from the number of detected genes and reads, percent mitochondrial genes and patient-specific differences were regressed out with NaiveDE v1.2.0 (https://github.com/Teichlab/NaiveDE) using the highly variable genes and retaining variation explained by antigen specificity. The expression matrix was scaled using sklearn (scikit-learn v1.0.2).^75^ UMAP plots were created with the *umap* function based on the first four principal components. The “MBC” and “PB” clusters were identified by unsupervised clustering using the *louvain* function. Differentially expressed genes between TG2-specific and other specificity cells were determined separately for the “MBC” and “PB” clusters with the *rank_genes_groups* function for genes expressed in at least 30% of the TG2-specific or other specificity groups, using Wilcoxon rank-sum test and the Benjamini-Hochberg method for correcting for multiple testing. Differentially expressed genes were visualized with EnhancedVolcano v1.18.0 in R 4.3.1 with adjusted p-value < 0.05 and log_2_ fold-change > 0.4 as cutoffs. Dotplots were drawn with the *dotplot* function (mean_only_expressed=True).

*Droplet-based scRNA-seq.* The droplet-based scRNA-seq gene expression analysis was performed using Seurat v4.3.0.1^73^ in R 4.3.1. Cells were categorized as CD27+, CD27-, TG2-specific and other specificity based on barcoded anti-CD27 and TG2 tetramer labelings, which were normalized by centered log-ratio transformation (see Fig. 3D). Differentially expressed genes between CD27+ and CD27- plasma cells were determined separately for TG2-specific and other specificities with the *rank_genes_groups* function for genes expressed in at least 30% of the TG2-specific or other specificity groups, using Wilcoxon rank-sum test and the Benjamini-Hochberg method for correcting for multiple testing. Differentially expressed genes were visualized with EnhancedVolcano v1.18.0 in R v4.3.1 with adjusted p-value < 0.05 and log_2_ fold-change > 0.3 as cutoffs.

### BCR reconstruction and clonal assignments

*Smart-seq2.* Paired BCR heavy- and light chains were reconstructed from the Smart-seq2 data using BraCeR.^76^ BraCeR assembly was run with *--threshold 500* to filter out lowly expressed reconstructed BCRs that may arise from contamination between wells or errors in indices. In addition to our Smart-seq2 data from blood, we retrieved BraCeR-reconstructed BCR sequences (EGA accession number EGAS00001004623) from our previous study containing Smart-seq2 data from small intestinal plasma cells of the same two patients.^3^ Only cells of the IgA isotype were retained. Clonally related cells were determined separately for each patient, antigen specificity (TG2 or other) and tissue type as well as for both tissue types together using BraCeR in summary mode. Potential cell multiplets were manually inspected as previously outlined.^77^

*Droplet-based scRNA-seq and Ig sequencing.* BCR sequences and clonally related cells from the droplet-based approach were identified by Cell Ranger for each patient (see **scRNA-seq quality control**). Only productively rearranged sequences belonging to cells passing quality control were retained. In rare cases where multiple heavy or light chains were reported for a cell, only the most highly expressed heavy and light chain were retained. The TG2-specific BCR sequences from droplet-based scRNA-seq and Ig sequences were additionally run through BraCeR in assemble mode and subsequently summary mode for each patient to identify clonally related cells between gut and blood. Clonal assignments were based on heavy chain (using DefineClones function of Change-O with CDR3 nucleotide distance < 0.2) due to the lack of light chain information for many of the gut PCs. Clone groups were manually inspected to verify that no clones had unrelated light chains when light chain sequences were present. Lineage trees based on heavy chain were also drawn separately for TG2-specific gut plasma cells through BraCeR.

### Mutational and repertoire analysis of BCRs

The BCR sequences from Smart-seq2, droplet-based scRNA-seq and Ig sequencing were analyzed using IMGT/HighV-QUEST^78^ in April 2023. Productive sequences from cells passing the quality control outlined above were parsed and aligned to germline sequences with the Change-O package (v.1.3.0)^79^ and analyzed for mutational load using Shazam v.1.1.2^79^. Repertoire usage as frequency of clones determined by BraCeR (Smart-seq2 and Ig sequencing) and Cell Ranger (droplet-based sequencing) was done using Alakazam v.1.2.1. Only patients with sequences from at least 50 TG2-specific and 50 other specificity blood cells were retained in the repertoire analysis.

### Statistical analysis

Statistical tests and calculation of p values were performed in GraphPad Prism v9 or ggpubr v0.6.0. Details about statistical tests, centers and error bars are given in the figure legends. P values below 0.05 were considered significant.

