## Supplemental figures 1-7 and Table S1 for "Generation of circulating autoreactive pre-plasma cells fueled by naïve B cells in celiac disease"

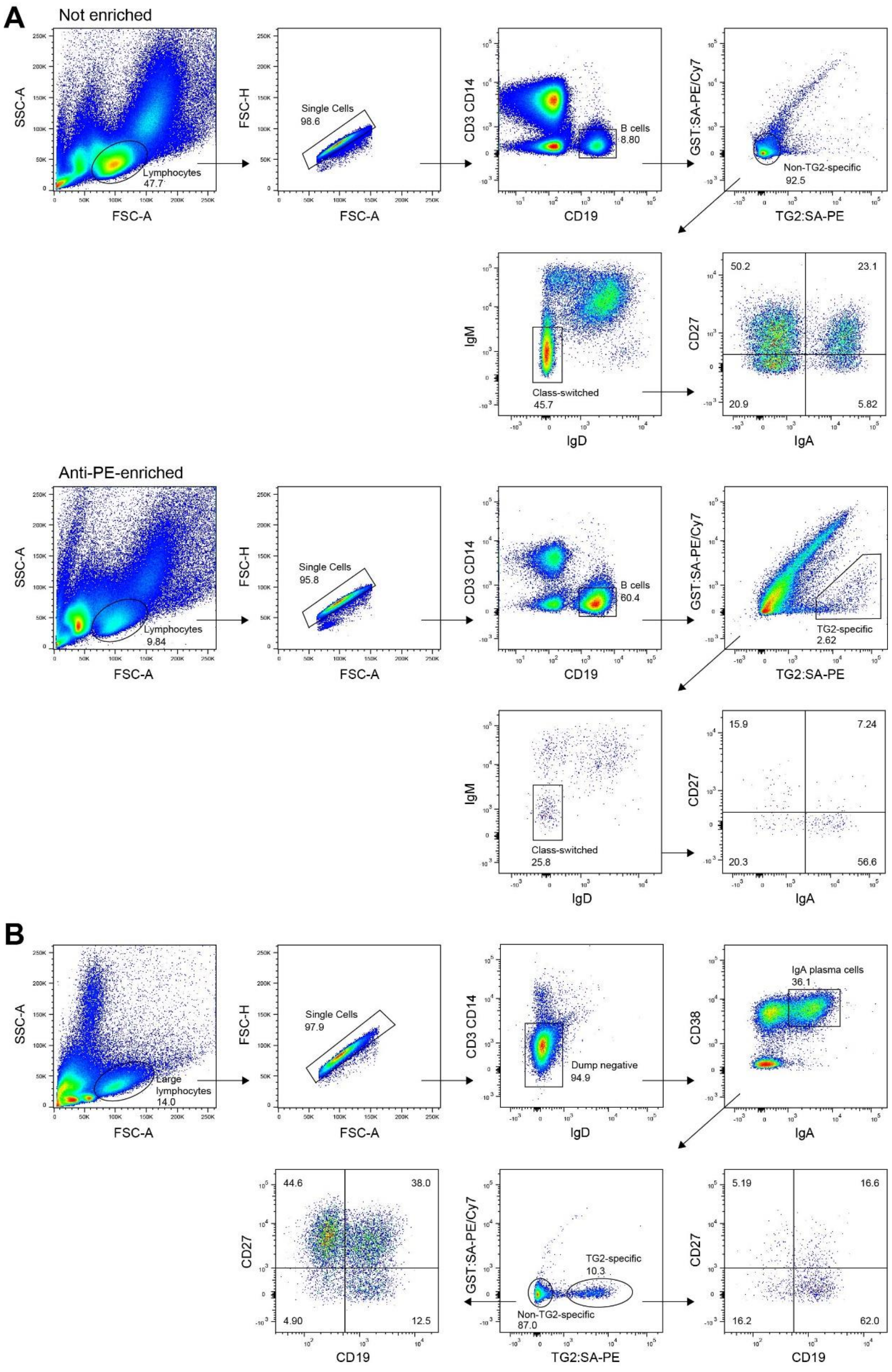


**Figure S1. Gating strategy.** (A and B) representative flow cytometry plots showing the general gating strategy used to identify TG2-specific and non-TG2-specific B cells and plasma cells in peripheral blood (A) and duodenal biopsy single-cell suspensions (B).

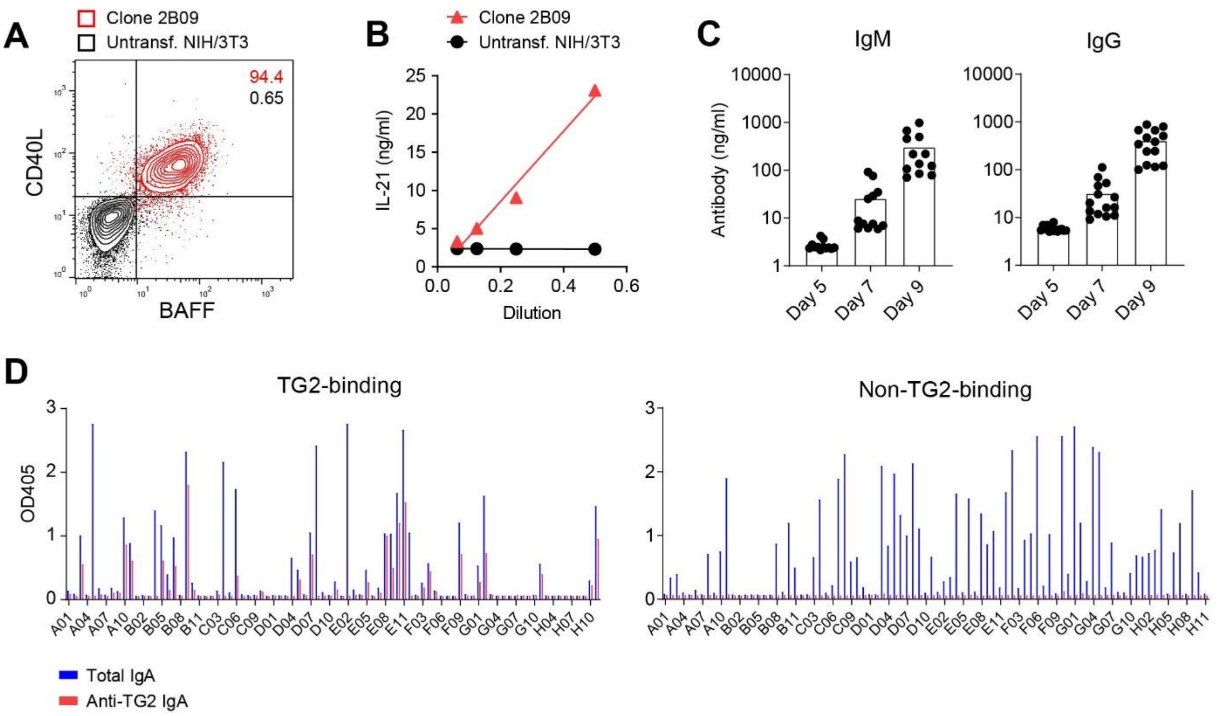


**Figure S2. Single-B-cell culture system.** (A and B) Verification of transgene expression by triple-transfected NIH/3T3 fibroblasts (clone 2B09). Surface expression of human BAFF and CD40L was confirmed by flow cytometry (A), and secretion of human IL-21 into the culture medium was assessed by ELISA (B). (C) Secretion of IgM and IgG into culture supernatants of single B cells isolated from PBMCs of a healthy donor. Total B cells were seeded into wells coated with 2B09 fibroblasts on day 0, and the amount of antibody secretion was determined at various time points by ELISA. (D) Detection of total and anti-TG2 IgA secretion in individual wells containing single TG2-binding or non-TG2-binding IgA+ B cells sorted from PBMCs of an untreated CeD patient. The cells were cultured for nine days together with 2B09 fibroblasts before antibody secretion was assessed by ELISA.

**A**

Sequencing-based isotype of IgA-sorted gut PCs passing quality control before filtering out IgM+ cells

| Donor | TG2-specific | | | Other | | |
|---|---|---|---|---|---|---|
| | IgA | IgM | % IgM | IgA | IgM | % IgM |
| CD1970 | 38 | 8 | 17.4 | 486 | 26 | 5.1 |
| CD2163 | 75 | 2 | 2.6 | 690 | 5 | 0.7 |
| CD5016 | 170 | 6 | 3.4 | 765 | 0 | 0.0 |
| CD5031 | 44 | 47 | 51.6 | 369 | 31 | 7.8 |

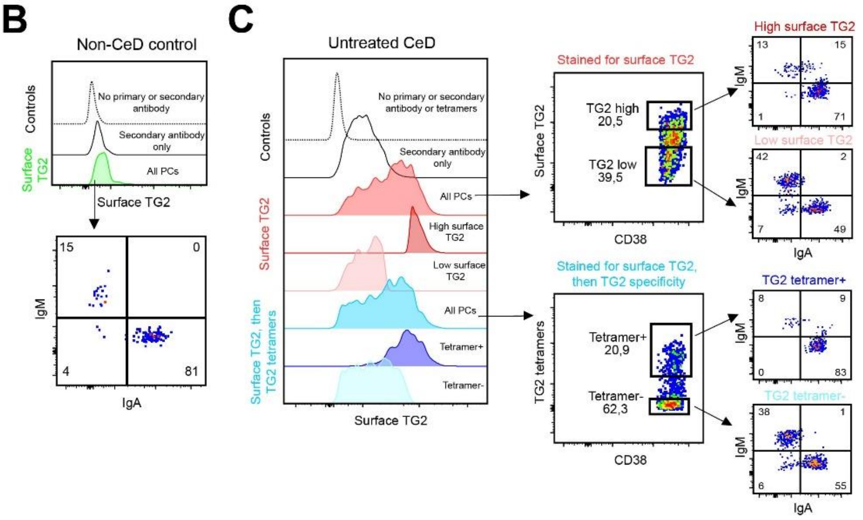


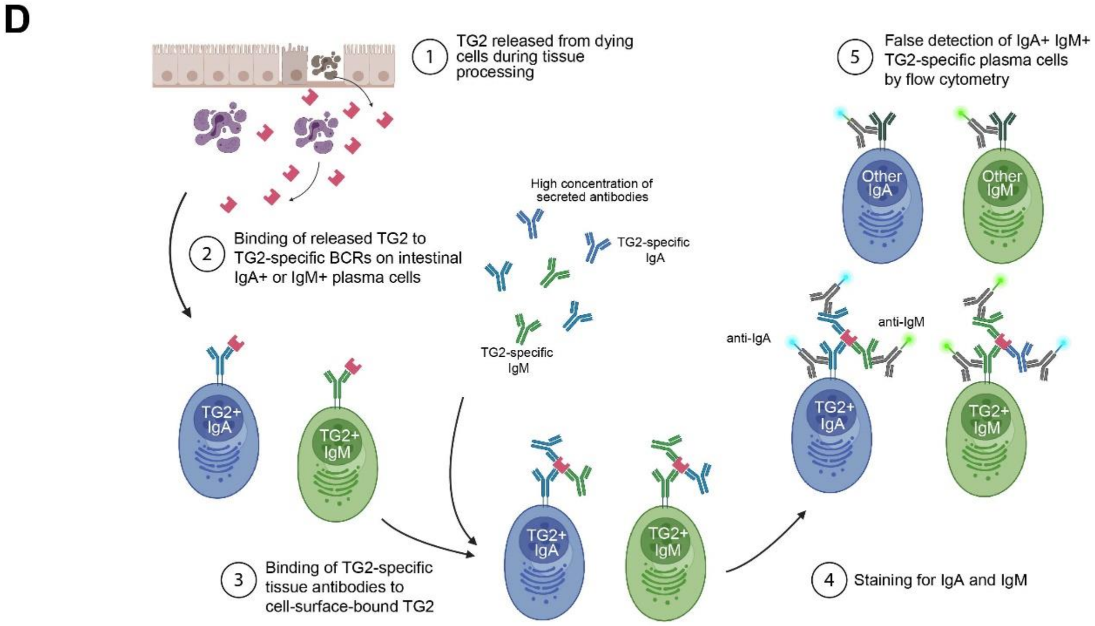


**Figure S3. IgA and IgM expression by TG2-specific gut plasma cells.** (A) Sequencing-based isotype of IgA-sorted gut plasma cells passing quality control, before removal of non-IgA cells. The isotype was derived from the V(D)J sequencing data generated by droplet-based sequencing (10x Genomics). A substantial number of IgA-sorted TG2-specific plasma cells in two of the donors turned out to be IgM+ based on the sequencing, illustrating that IgA staining of TG2-specific cells may not always represent the true isotype of the cells. (B and C). Single cell suspensions from small intestinal biopsies of healthy controls (B, n=2) or untreated CeD patients (C, n=2) were stained for IgA, IgM and presence of surface-bound TG2 using a rabbit anti-human TG2 primary antibody and an anti-rabbit IgG secondary antibody. Representative data from one of two independent experiments are shown. Plasma cells were identified as large $CD3^{-}$ $CD11c^{-}$ $CD14^{-}$ $CD38^{++}$ cells. (C) Samples from the CeD patients were additionally stained for TG2-specific cells using streptavidin TG2 tetramers (n=1) or biotinylated TG2 monomers followed by detection with an anti-biotin antibody (n=1). (D) Proposed mechanism underlying increased numbers of TG2-specific plasma cells staining positive for both IgA and IgM simultaneously compared to cells of other specificities or healthy controls.

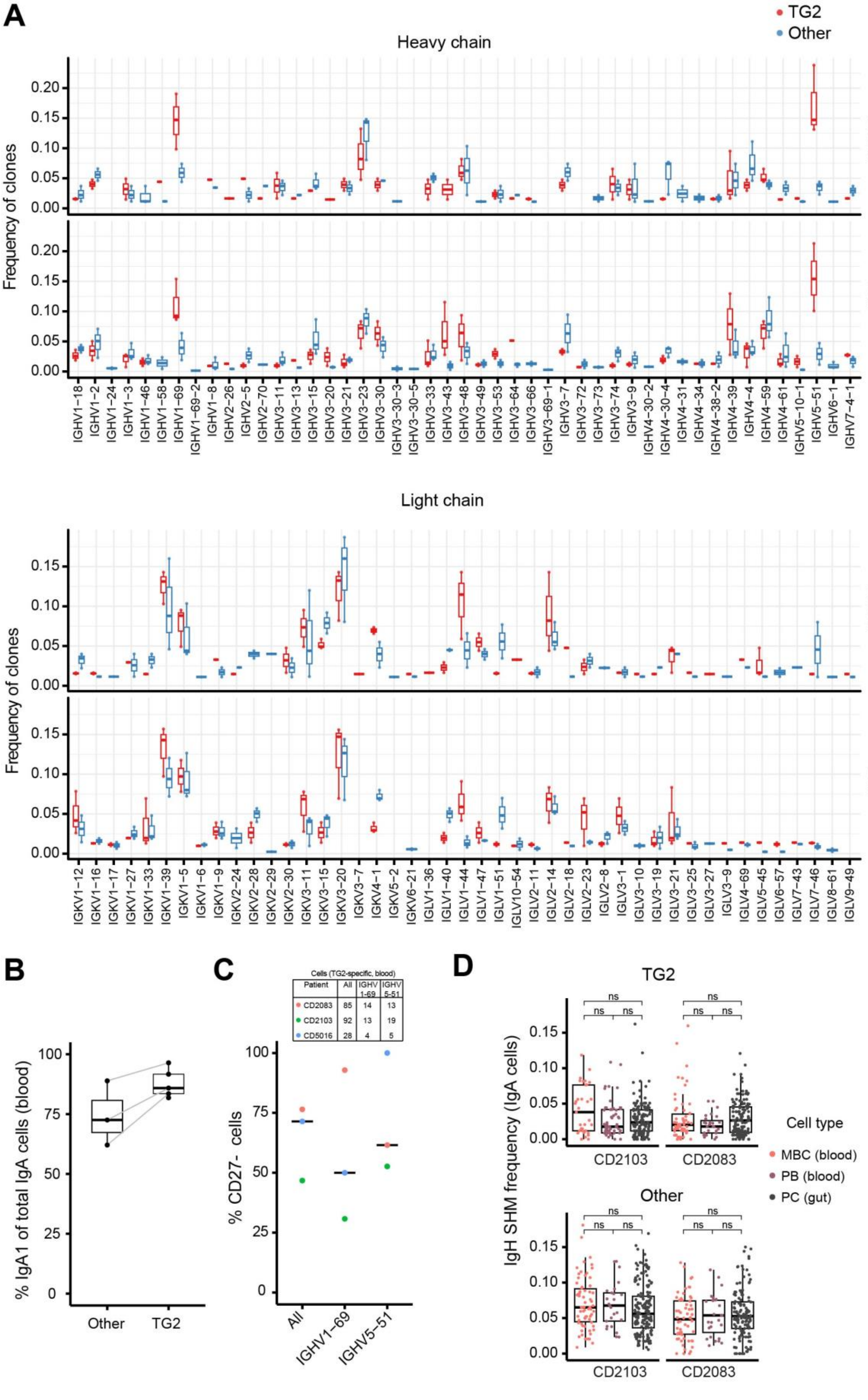


| Patient | All | IGHV 1-69 | IGHV 5-51 |
|---|---|---|---|
| CD2083 | 85 | 14 | 13 |
| CD2103 | 92 | 13 | 19 |
| CD5016 | 28 | 4 | 5 |

**Figure S4. Ig sequence properties of TG2-specific IgA cells.** (A) Frequency of individual IGHV and IGKV/IGLV gene segments among TG2-specific and non-TG2-specific (other) IgA clones in blood and gut biopsies of untreated CeD patients (n=3). Boxes indicate the first and third quartiles; the horizontal line indicates median value. Only patients where at least one clone used an indicated V-gene is included in each plot. (B) Proportion of circulating TG2-specific and other IgA cells expressing the IgA1 subclass in individual patients with untreated CeD. (C) Frequency of CD27- cells among all TG2-specific IgA cells and those using two prominent IGHV segments in blood of three untreated CeD patients. (D) Comparison of mutation levels between IgA memory B cells (MBC), plasmablasts (PB) and plasma cells (PC) in two untreated CeD patients. Centers indicate medians, and statistical significance was evaluated by a Mann-Whitney U test.

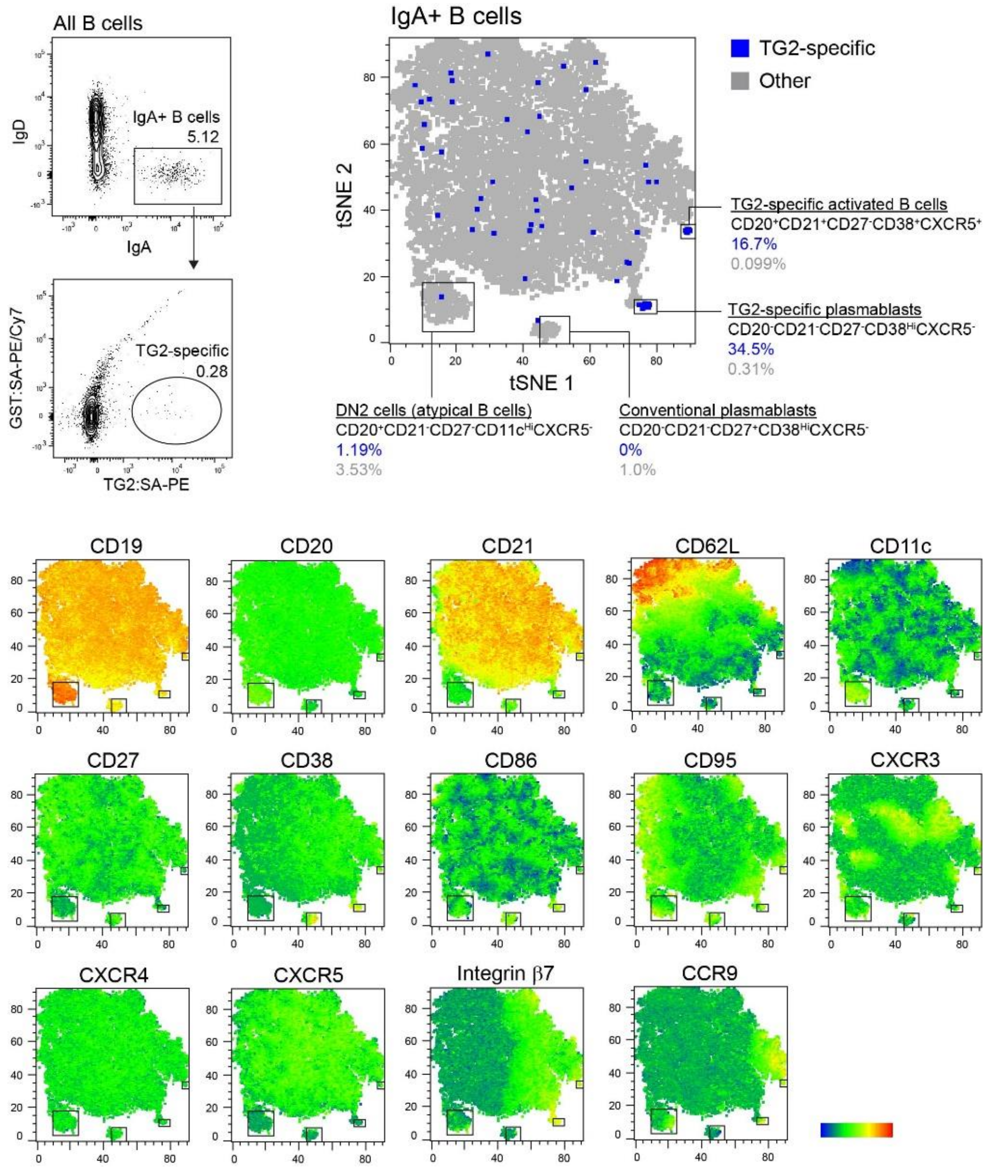


**Figure S5. Phenotypic characterization of circulating TG2-specific B cells by flow cytometry.** Representative plots showing identification of TG2-specific and non-TG2-specific (other) IgA+ B cells in PBMCs of an untreated CeD patient without antigen-specific enrichment. The tSNE plot was generated from the 14 individual markers shown in the lower panels, and the gates were set based on markers associated with plasmablasts and classical or atypical memory B cells.

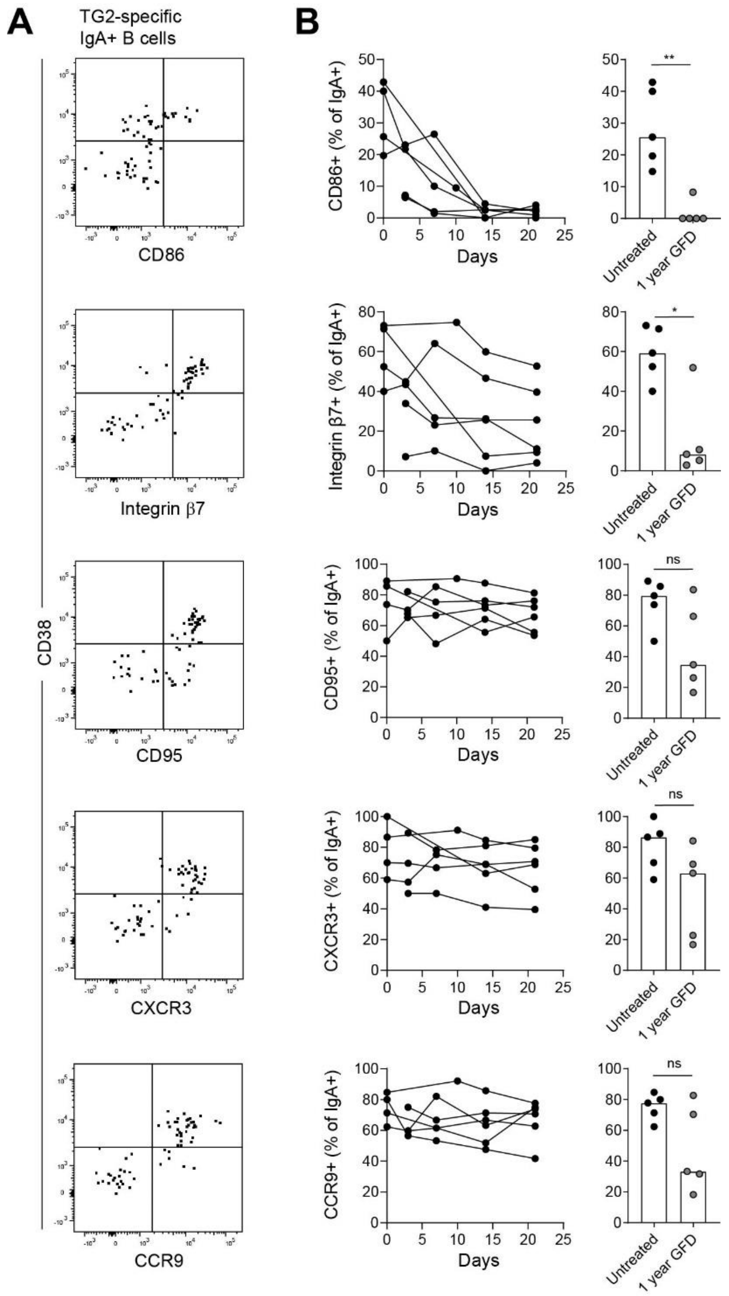


**Figure S6. Expression of surface markers before and after exclusion of dietary gluten.** (A) Representative flow cytometry plots showing gating of surface markers on TG2-specific IgA+ B cells in peripheral blood of an untreated CeD patient. (B) Change in phenotype of TG2-specific IgA+ B cells in individual patients (n=6) during the first three weeks of gluten-free diet (GFD) and comparison of untreated patients and patients who had been on GFD for one year. Bar heights indicate medians, and difference between groups was evaluated by a Mann-Whitney U test. *p < 0.05, **p < 0.01.

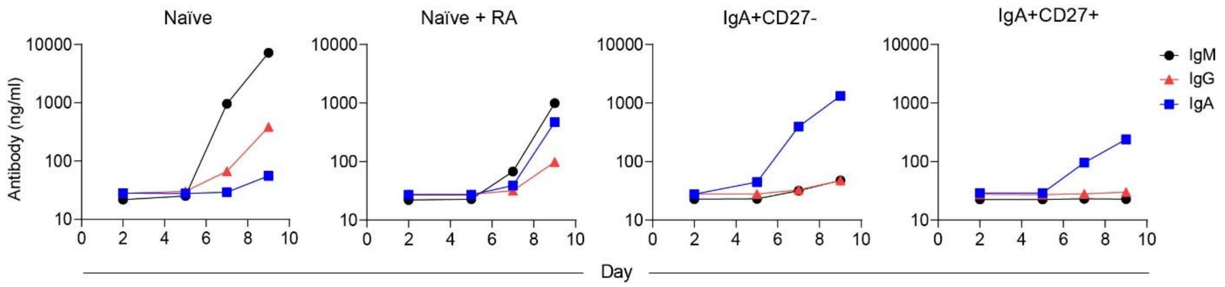


**Figure S7. Antibody secretion in B-cell cultures.** Detection of total IgM, IgG and IgA in cultures of different B-cell populations sorted from peripheral blood of a healthy donor. The B cells were placed in culture with transfected fibroblasts on day 0, and antibody secretion was assessed at various time points by ELISA. Naïve (IgD+CD27-) B cells were cultured with or without addition of retinoic acid (RA) on day 0.

**Table S1. Clinicopathological data of untreated CeD patients used for sequencing analyses of peripheral blood B cells and duodenal plasma cells.**

| | Patient ID | Gender | Age | Anti-TG2 IgA[1] (cut-off 5) | Anti-DGP IgG[1] (cut-off 20) | Marsh score[2] |
|---|---|---|---|---|---|---|
| Ig sequencing of single-B-cell cultures and transcriptome of gut plasma cells (10x Genomics) | CD1970 | M | 64 | >100 | 13 | 3B |
| | CD2163 | F | 80 | >100 | >100 | 3B |
| | CD5016 | F | 46 | 32[3] | 20[3] | 3A |
| | CD5031 | F | 26 | >128[3] | 242[3] | 3B-C |
| Transcriptome of blood B cells (Smart-seq2)[4] | CD2083 | M | 29 | >100 | >100 | 3C |
| | CD2103 | F | 32 | 11.7 | >100 | 3C |

[1]Serum levels of TG2-specific IgA and deamidated gluten peptide (DGP)-specific IgG at the time of examination.
[2](Oberhuber et al., 1999).
[3]Cut-off 7.
[4]Transcriptomic data of gut plasma cells isolated from these two patients are reported in (Lindeman et al., 2021).
